# Generation of an inflammatory niche in an injectable hydrogel depot through recruitment of key immune cells improves efficacy of mRNA vaccines

**DOI:** 10.1101/2024.07.05.602305

**Authors:** Emily L. Meany, John H. Klich, Carolyn K. Jons, Tianyang Mao, Namit Chaudhary, Ashley Utz, Julie Baillet, Ye E. Song, Olivia M. Saouaf, Ben S. Ou, Shoshana C. Williams, Noah Eckman, Darrell J. Irvine, Eric Appel

## Abstract

Messenger RNA (mRNA) delivered in lipid nanoparticles (LNPs) rose to the forefront of vaccine candidates during the COVID-19 pandemic due in part to scalability, adaptability, and potency. Yet there remain critical areas for improvements of these vaccines in durability and breadth of humoral responses. In this work, we explore a modular strategy to target mRNA/LNPs to antigen presenting cells with an injectable polymer-nanoparticle (PNP) hydrogel depot technology which recruits key immune cells and forms an immunological niche in vivo. We characterize this niche on a single cell level and find it is highly tunable through incorporation of adjuvants like MPLAs and 3M-052. Delivering commercially available SARS-CoV-2 mRNA vaccines in PNP hydrogels improves the durability and quality of germinal center reactions, and the magnitude, breadth, and durability of humoral responses. The tunable immune niche formed within PNP hydrogels effectively skews immune responses based on encapsulated adjuvants, creating opportunities to precisely modulate mRNA/LNP vaccines for various indications from infectious diseases to cancers.

## 1. Introduction

Over the last three decades, messenger RNA (mRNA) technology has grown as a therapeutic modality for treatment and prevention of various diseases. It has been developed for protein replacement as well as vaccination against infectious diseases and cancers (*1–3*). During the COVID-19 pandemic, mRNA delivered in lipid nanoparticles (LNPs) rose to the forefront of vaccine candidates due to its safety profile, rapid and affordable scalability, and adaptability to emergent variants of concern. Moderna and Pfizer-BioNTech mRNA vaccines proved highly potent and were instrumental in curbing the spread of SARS-CoV-2. Yet there remain critical areas for improvement of mRNA vaccines, including stability for simpler storage, transport, and deployment in under-resourced parts of the world, as well as enhanced durability and breadth of humoral immune responses (*4–6*). In this work, we develop an approach to enhancing mRNA/LNP vaccines by targeting of the mRNA/LNPs to antigen presenting cells (APCs), adjuvanting with molecular adjuvants, and prolonging antigen availability.

APCs are key immune cells responsible for processing and presenting antigen to train the adaptive arm of the immune system—T and B cells—and a desirable target for mRNA/LNP transfection (*7*). A recent study by Hassett et al. at Moderna found macrophages and dendritic cells (DCs) infiltrating the site of administration, as well as off-target adipocytes, fibroblasts, and epithelial cells in these tissues, took up mRNA and expressed protein following intramuscular (IM) injection in non-human primates (NHPs), with peak expression 24 hours post-injection (*8*). While antigen produced by non-APCs could contribute to a meaningful immune response, improving antigen production and delivery to professional APCs is expected to be beneficial for priming both B and T cell responses. Many groups have worked to improve APC targeting by modifying LNPs with targeting moieties like antibodies or mannose, as well as altering lipid chemistries to bias their biodistribution (*9–12*). While many of these strategies have demonstrated improved cell-type targeting, they often involve cumbersome large-scale screening campaigns and complex lipid chemistries that can alter LNP bioactivity since the lipids are known to contribute to transfection efficiency and adjuvanting effects of the LNPs (*13, 14*). Moreover, each newly designed lipid will be regulated as a novel chemical entity, thereby limiting translatability. mRNA/LNP vaccines could benefit from a more modular method to target APCs and limit off-target cell transfection without alterations to the existing LNP chemistries.

In addition to better APC targeting, adjuvanting mRNA/LNP vaccines could improve the overall magnitude and quality of the vaccine response. Adjuvants are immunostimulants which can drive potent immune responses, as well as tailor a response toward T helper-1 (Th1), Th2, or other phenotypes important in mounting successful responses against different diseases. Yet the impact of adjuvants on mRNA vaccines remains poorly characterized, with evidence of both benefits and detriments previously reported (*15–17*). Excessive stimulation of innate cells can reduce mRNA expression and negatively impact vaccine efficacy. Seemingly contrary to this, some groups have found incorporation of pathogen-associated molecular pattern (PAMP) adjuvants like toll-like receptor agonists (TLRas) can improve mRNA vaccine efficacy, particularly TLR2/6a, 7/8a, and 9a (*18–21*). While promising, these studies required lipid modification, reformulation, or even use of a different delivery modality entirely, and are not readily applicable within existing mRNA/LNP vaccines. Furthermore, the need to modify LNPs to incorporate adjuvants severely limits which adjuvants can be used and compromises the ability to directly compare impacts of adjuvants across platforms. An off-the-shelf technique allowing modular selection of single or combination adjuvants will be especially beneficial to cater to different disease needs as mRNA vaccines expand in use to other infectious diseases, anti-cancer vaccines, and even tolerogenic vaccines (*22*).

Finally, it has been shown that extended delivery of antigen can improve germinal center reactions and overall vaccine responses to subunit vaccines (*23–25*). However, sustained delivery is difficult to achieve with mRNA vaccines as mRNA expression is transient and mRNA is prone to hydrolysis and degradation following injection, even with advances in base modifications and LNP delivery vehicles. Previous efforts have pursued prolonged antigen availability through increased expression with self-amplifying RNA (saRNA), but LNPs for saRNA can be difficult to formulate due to the larger mRNA required to encode the target antigen as well as the replication machinery (*26*). A means to extend the expression and availability of antigen with standard mRNA vaccines could further improve their efficacy.

In this work, we explored a modular strategy to adjuvant mRNA/LNPs and target their uptake by APCs via encapsulation in an injectable, dynamic hydrogel depot technology known to attract and activate key immune cells in vivo in a local immunologic niche. We leveraged polymer-nanoparticle (PNP) hydrogels formed through dynamic, non-covalent interactions between hydrophobically-modified hydroxypropylmethylcellulose (HPMC-C_12_) and poly(ethylene glycol)-block-poly(lactic acid) nanoparticles (PEG-PLA NPs) (*27*). These materials are biocompatible and inert, yet they attract immune cells and form an immune niche in vivo when loaded with inflammatory cargo like vaccines and adjuvants (*28–30*). While these hydrogels have a small effective mesh size capable of entrapping diverse molecular cargo, their dynamic crosslinks allow cells to exert forces and actively migrate into the material and interact with loaded cargos (*31–37*). We hypothesized these materials would promote targeting of mRNA/LNPs to infiltrating APCs, as well as allow for admixing of different adjuvants for a modular, off-the-shelf platform to study and compare adjuvanting of mRNA vaccines. The ability to admix different adjuvants allows for tuning of the inflammatory microenvironment where APCs encounter the mRNA/LNPs across a spectrum of inflammatory states, from highly inflammatory for anti-cancer applications to tolerogenic for applications in diabetic “inverse” vaccines. We also hypothesized that sequestering mRNA/LNPs within the hydrogel would slow the diffusion and penetration of RNases and other enzymes, allowing for extended delivery and expression of mRNA, improving the immune response.

We first demonstrated the plasticity of the PNP hydrogel immunologic niche in vivo in response to mRNA/LNPs alone or with two separate TLRas, synthetic monophosphoryl lipid A (synthetic MPLA, denoted MPLAs; TLR4a) or 3M-052 (TLR7/8a). Flow cytometry analyses revealed clear skewing of the cellular milieu based on encapsulated cargo, with transient infiltration of different immune cells such as neutrophils, monocytes, or natural killer cells influenced by different cargos. Using a model mRNA cargo, we examined the mRNA expression in this in vivo niche and found key APCs actively take up LNPs and produce the encoded protein. We investigated the impact of distinct PNP hydrogel-based immune niches on the humoral and cellular responses to a commercially available SARS-CoV-2 mRNA vaccine compared with a clinical control bolus administration. Both PNP hydrogels alone and with co-encapsulated adjuvants improved the magnitude, durability, and breadth of humoral immune responses, the functionality of the antibodies produced, and the systemic T cell response. We further investigated the lymph node germinal center reactions underlying these improvements in vaccine response and found PNP hydrogels, particularly when adjuvanted, increased the magnitude and quality of the germinal center reaction to mRNA/LNP vaccines.

## 2. Results

### 2.1 Characterization and in vitro validation of supramolecular hydrogels loaded with mRNA/LNPs

In this work, we leveraged the unique mechanical properties and modularity of PNP hydrogels to form an immune niche and selectively deliver mRNA/LNPs to recruited APCs. Following subcutaneous injection in mice, migratory immune cells are recruited to the injected hydrogel depot, where they can exert forces and infiltrate the dynamically crosslinked material (**Figure 1A**). Once cells enter the PNP hydrogel depot, they encounter the mRNA/LNPs and can be transfected before migrating to the draining lymph node to initiate germinal center reactions. We hypothesized we could skew the cell populations in the hydrogel immune niche by incorporating different inflammatory cargo such as potent TLR4a (MPLAs) or TLR7/8a (3M-052) adjuvants. These potent adjuvants have shown promise in other vaccine formulations and clinical vaccines (e.g., the human papillomavirus vaccine by GSK is adjuvanted with ASO4 comprising MPL/Alum) (*38–40*). Additionally, both adjuvants have lipid moieties which promote retention in the PNP hydrogel (*37*).

**Fig. 1.**
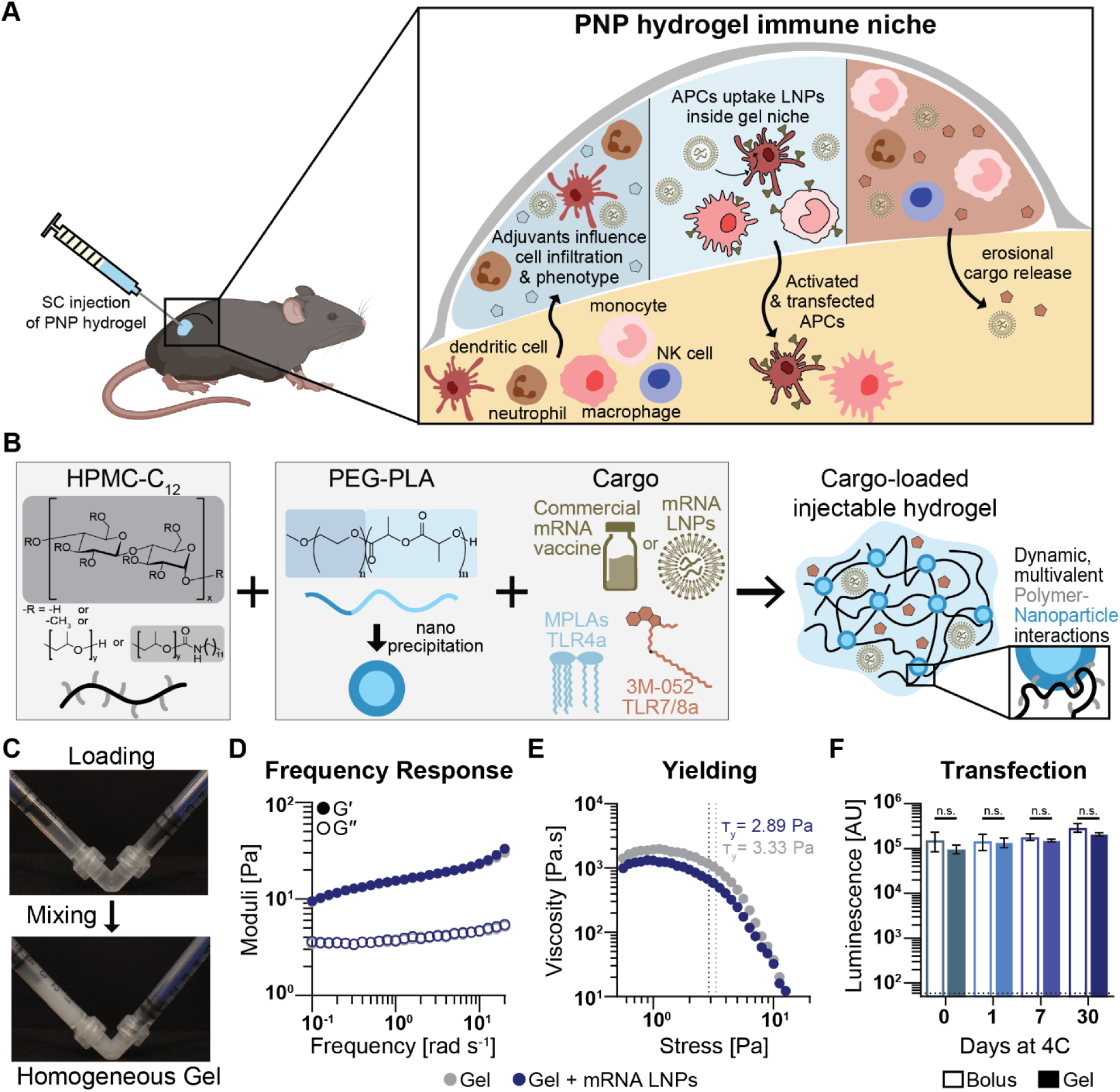
Schematic of immune niche in PNP hydrogels comprising mRNA/LNPs. A) PNP hydrogels injected subcutaneously allow immune cell infiltration and interaction with loaded cargo. Incorporation of various adjuvants influences the immune niche. Cells within the niche take up mRNA/LNPs, express the delivered protein, and migrate to initiate immune responses in the draining lymph nodes. B) PNP hydrogels comprise HPMC-C_12_ and PEG-PLA NPs. Cargo, including mRNA/LNP vaccines and molecular adjuvants, can be easily mixed with the polymer components to form a dynamic hydrogel. C) Hydrogel components are loaded into two syringes, attached with an elbow, and mixed to form a homogenous material, pre-loaded in a syringe and ready for administration. D) Frequency shear rheology shows PNP-0.5-5 material is solid-like across time scales and mRNA/LNPs (50 μg mL^-1^) do not alter this. E) Stress-ramp data shows static yield stress, defined as the intersection of tangent lines for the plateau and yielding regimes, for PNP-0.5-5 hydrogel is not impacted by LNPs. F) Luminescent signal from RAW-Blue macrophages dosed with luciferase mRNA/LNPs in either PBS bolus or PNP hydrogel after storage in that condition for up to 30 days at 4 °C. Hydrogels do not impact transfection or stability of mRNA/LNPs. Data shown as mean ± SD, n = 3 - 4, and statistics are multiple unpaired two-tailed t-tests run in GraphPad Prism with false discovery rate (FDR) correction using two-stage step-up method of Benjamini, Krieger, and Yekutieli.

PNP hydrogels are formed through simple mixing of HPMC-C_12_ and PEG-PLA NPs. These components interact in a multivalent, non-covalent fashion to form an injectable shear-thinning and self-healing dynamic hydrogel. Multiple cargos can be admixed into the hydrogel, including commercially available mRNA vaccines, other mRNA/LNPs, and adjuvants (**Figure 1B-C**). We first sought to characterize how commercially available SARS-CoV-2 mRNA vaccine impacts key mechanical properties of the PNP hydrogels. Frequency sweep oscillatory shear rheology showed the selected formulation, PNP-0.5-5 (0.5 wt% HPMC-C_12_; 5 wt% NPs), was solid-like over tested frequencies and was not impacted by mRNA/LNP incorporation (**Figure 1D**). Further, stress ramp experiments showed clear pre-yielded and yielded regimes with a yield stress of approximately 3 Pa (**Figure 1E**). These results indicated these materials would form disk-like depots in the subcutaneous space (*41*). High-to-low shear-ramp and step-shear experiments showed that PNP-0.5-5 hydrogels were shear-thinning and capable of repeatedly returning to a high viscosity state following high-shear events, indicating the hydrogel could be injected through standard needles and recover as a solid-like depot following injection (**Supp Fig. S1**).

Having shown mRNA/LNPs did not disrupt hydrogel material properties, we next confirmed that mRNA/LNPs were unimpacted by loading within PNP hydrogels. We assessed mRNA/LNP transfection efficiency in vitro with luciferase-encoding mRNA/LNPs mimicking the lipid formulation used in Moderna’s Spikevax vaccine. In these assays, we diluted the luciferase mRNA/LNPs into PBS or incorporated them into PNP-0.5-5 hydrogels and stored at 4 °C for 0, 1, 7, and 30 days before delivering to RawBlue macrophages and measuring transfection efficiency via bioluminescence after 24 hours (**Figure 1F**). Transfection efficiency remained unchanged for mRNA/LNPs formulated into PNP hydrogels compared with delivery in bolus at all tested storage times. These stability data agreed with the product stability for Moderna Spikevax (4 °C for 30 days) and demonstrated PNP hydrogel-based formulations did not impede transfection capacity of the mRNA/LNPs.

### 2.2 Tunable immune niche formed within PNP hydrogel in vivo

One unique feature of PNP hydrogels is that their nano-scale mesh size allows retention of small cargos, yet their dynamic crosslinks allow immune cells to migrate into the material in response to inflammatory cargo (*42*). We employed flow cytometry to phenotype the immune niche within the PNP hydrogel depot over time in response to mRNA/LNPs with and without MPLAs or 3M-052 adjuvants. We injected C57BL/6 mice subcutaneously on the flank with 100 μL of PNP hydrogel containing 1 μg mCherry mRNA/LNPs, either alone or with adjuvants, and excised the hydrogel depots at days three and seven for single cell flow cytometry (**Figure 2A, Supp Fig. S2**). The hydrogels clearly formed persistent depots without signs of fibrosis or foreign body response on both excision days (**Figure 2B, Supp Fig. S3**). The hydrogels contained over a million CD45^+^ leukocytes on day three (1.35 M), and more in adjuvant-loaded hydrogels (2.23 M with MPLAs and 2.27 M with 3M-052; *p* = 0.071 for MPLAs and 0.057 for 3M-052 compared to LNP-only hydrogels). PNP hydrogels with mRNA/LNPs hosted almost double the cells found in empty PNP hydrogels (0.78 M), likely due to the inherent immunogenicity of mRNA/LNPs, and PNP hydrogels loaded with Moderna Spikevax showed comparable infiltration to ones loaded with mCherry mRNA/LNPs (**Supp Fig. S4**). Over time, cells migrate out of PNP hydrogels and the depot dissolves away. On day seven we found fewer, but still a measurable number of, cells in LNP-only hydrogels (0.20 M), while adjuvanted hydrogels maintained more cells (0.34 M with MPLAs and 0.65 M with 3M-052) (**Figure 2C**). The hydrogel depots measured approximately 100 mm^3^ on day three and 70 – 75 mm^3^ on day seven, indicating a cell density of 3–23 M mL^-1^. For reference, the average cell density in lymph nodes across animals studied for germinal center responses later in this work was ∼110 M mL^-1^ (*43*). Impressively, the PNP hydrogel depots sustained cell densities up to a quarter of that of lymph nodes undergoing active vaccine responses.

**Fig. 2.**
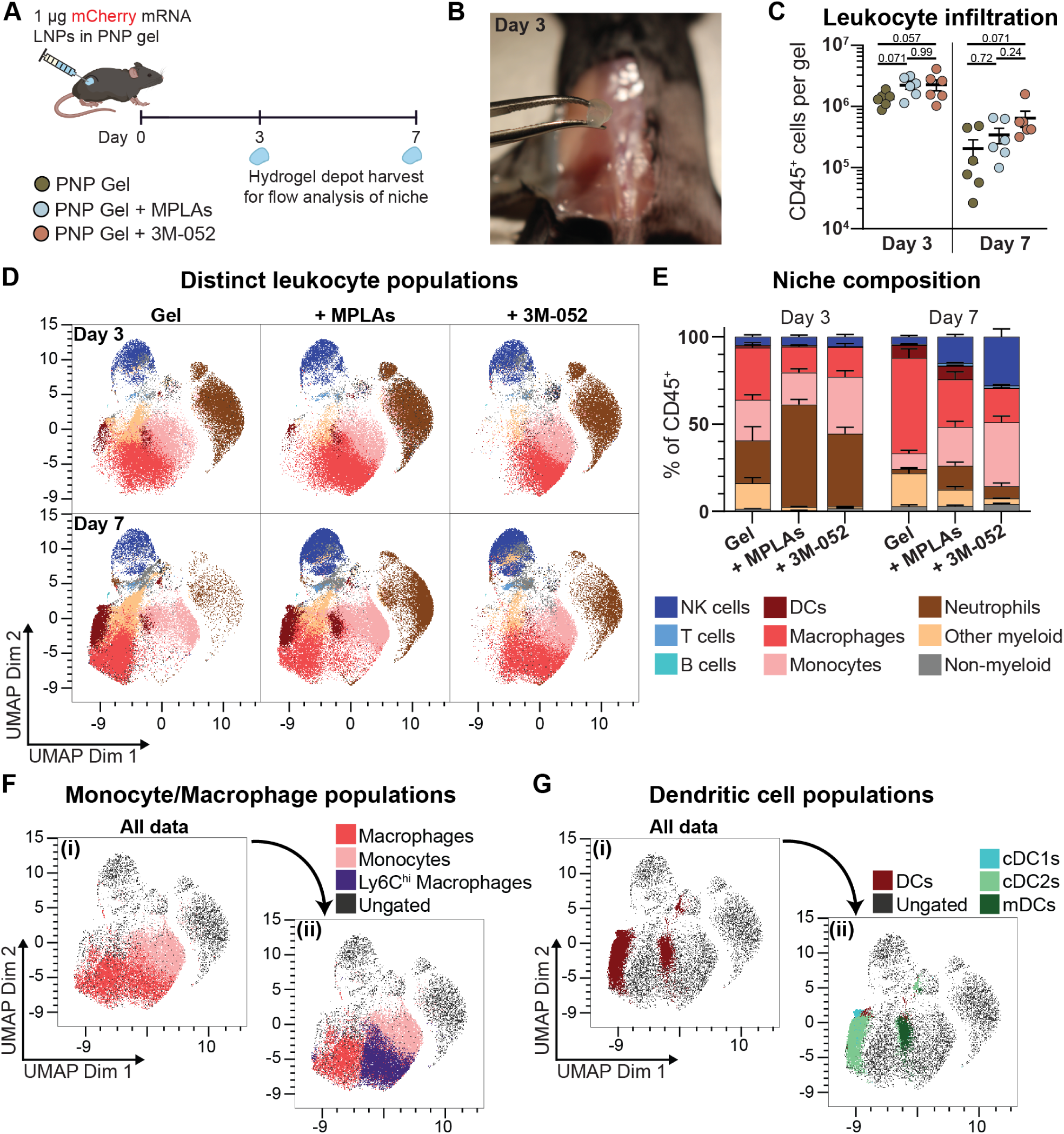
Characterization of in vivo immune niche in PNP hydrogels. A) PNP hydrogels loaded with mCherry mRNA/LNPs injected subcutaneously were excised at days 3 and 7 for dissociation and analysis of cells with flow cytometry. B) PNP hydrogel forms an excisable depot with no evidence of fibrosis or foreign body response on day 3. C) Counts of CD45^+^ leukocytes per gel show robust infiltration on both days, with increased infiltration with adjuvant cargo. Data shown as mean ± SEM, n = 6. Statistical values shown are *p* values obtained from general linear model (GLM) fitting and Tukey HSD multiple comparison test in JMP. D) Changes in cell type populations visualized with UMAPs. E) Quantification of cell types in each hydrogel niche as percentages of all gated CD45^+^ cells. Data shown as mean + SEM, n = 6. F) UMAP visualization showing relation of Ly6C^hi^ macrophage subpopulation to macrophages and monocytes. G) UMAP visualization of dendritic cell (DC) subpopulations highlighting differences between classical DCs (cDC1s and 2s) and monocyte-derived DCs (mDCs).

We then sought to examine the plasticity of the PNP cell niche across time and with the different adjuvant cargos using uniform manifold approximation and projection (UMAP) (**Figure 2D**). Overlaying the generated UMAP with gated cell populations, we found cell types fell within clear UMAP clusters and observed distinct and dramatic differences among groups and time points. We quantified these differences with the percentage for each cell type of the total CD45^+^ cells, as well as cell counts per gel (**Figure 2E, Supp Fig. S5**). On day three, there were clear differences across groups in neutrophil, DC, and monocyte populations. Neutrophils composed 59% and 42% of CD45^+^ cells in the MPLAs and 3M-052 groups, but only 24% in LNP-only hydrogels. There were more monocytes in adjuvanted groups compared to LNP-only hydrogels, significantly so for 3M-052 hydrogels (*p*=0.01). Surprisingly, fewer DCs were observed in 3M-052 hydrogels compared to LNP-only and MPLAs hydrogels (*p*=0.021 and 0.066 respectively). We also visualized different subpopulations and phenotypes of macrophages and DCs on the UMAPs, with monocyte-derived macrophages (Ly6C^hi^) and DCs (mDCs) clustering nearer to monocytes (**Figure 2F-G**). By day seven, all cell types in LNP-only hydrogels had decreased, with neutrophils composing 2% and macrophages 55% of all CD45^+^ cells. While macrophage counts in all PNP hydrogel groups were similar, the percentage these cells comprised of CD45^+^ cells, and their phenotypes, were distinct. Macrophages made up 27% of MPLAs hydrogels and spanned the UMAP spectrum, while they made up 19% of 3M-052 hydrogels and fell almost entirely within the monocyte-derived region of the UMAP. In contrast, LNP-only macrophages were 55% of the CD45^+^ cells and primarily Ly6C^lo^. Adjuvanted hydrogels retained substantially more neutrophils and monocytes at day seven (*p*=0.085 and *p*=0.024 for each cell type respectively in 3M-052 hydrogels). MPLAs hydrogels exhibited a slight increase in DCs with a larger population of inflammatory mDCs compared to LNP-only hydrogels. Finally, we also observed an influx of natural killer (NK cells) into adjuvanted groups, especially 3M-052 hydrogels, where they were found to comprise 28% of CD45^+^ cells. Overall, these data clearly indicate that immune cells migrate to injected PNP hydrogel depots and form highly plastic and tunable inflammatory niches in response to different adjuvant cargos. The ability of PNP hydrogel materials to retain vaccine cargo while enabling cellular infiltration is one which we can leverage to target mRNA/LNPs to key APCs under specific conditions, such as being surrounded by unique cellular milieu whose inflammatory state can be skewed with different cargos.

### 2.3 Expression pattern of delivered mRNA in PNP hydrogel cell niche

Following characterization of the highly plastic cell niche within the PNP hydrogel depot, we wanted to understand if cells within the niche took up LNPs and expressed the delivered mCherry mRNA (**Figure 3A**). We found that cells isolated from hydrogels on days three and seven were mCherry^+^ by flow cytometry (**Figure 3B**). We quantified the count and proportion of CD45^+^ mCherry^+^ cells and observed 3.8, 2.2, and 6.9 (×10^3^) cells expressed mCherry on day three in LNP-only, MPLAs, and 3M-052 hydrogels respectively, making up 0.3, 0.1, and 0.3% of all CD45^+^ cells. The PNP hydrogels clearly prolonged mRNA expression through day seven, with significantly more mCherry^+^ cells in adjuvanted groups compared to the LNP-only group, up to 12.3 (×10^3^) in 3M-052 hydrogels (2.1% of all CD45^+^ cells, *p* = 0.0008 and 0.0002 compared to LNP-only by count and percentage respectively) (**Figure 3 C, D**). Further examining the cell subtypes expressing delivered mRNA, we observed that mCherry^+^ cells were overwhelmingly APCs – DCs, monocytes, macrophages – across all gel groups (**Figure 3E, Supp Fig. S6**). Interestingly, monocytes composed 48% and 76% of mCherry^+^ cells in MPLAs and 3M-052 hydrogels, whereas DCs composed 22% and only 1% for each group respectively.

**Fig. 3.**
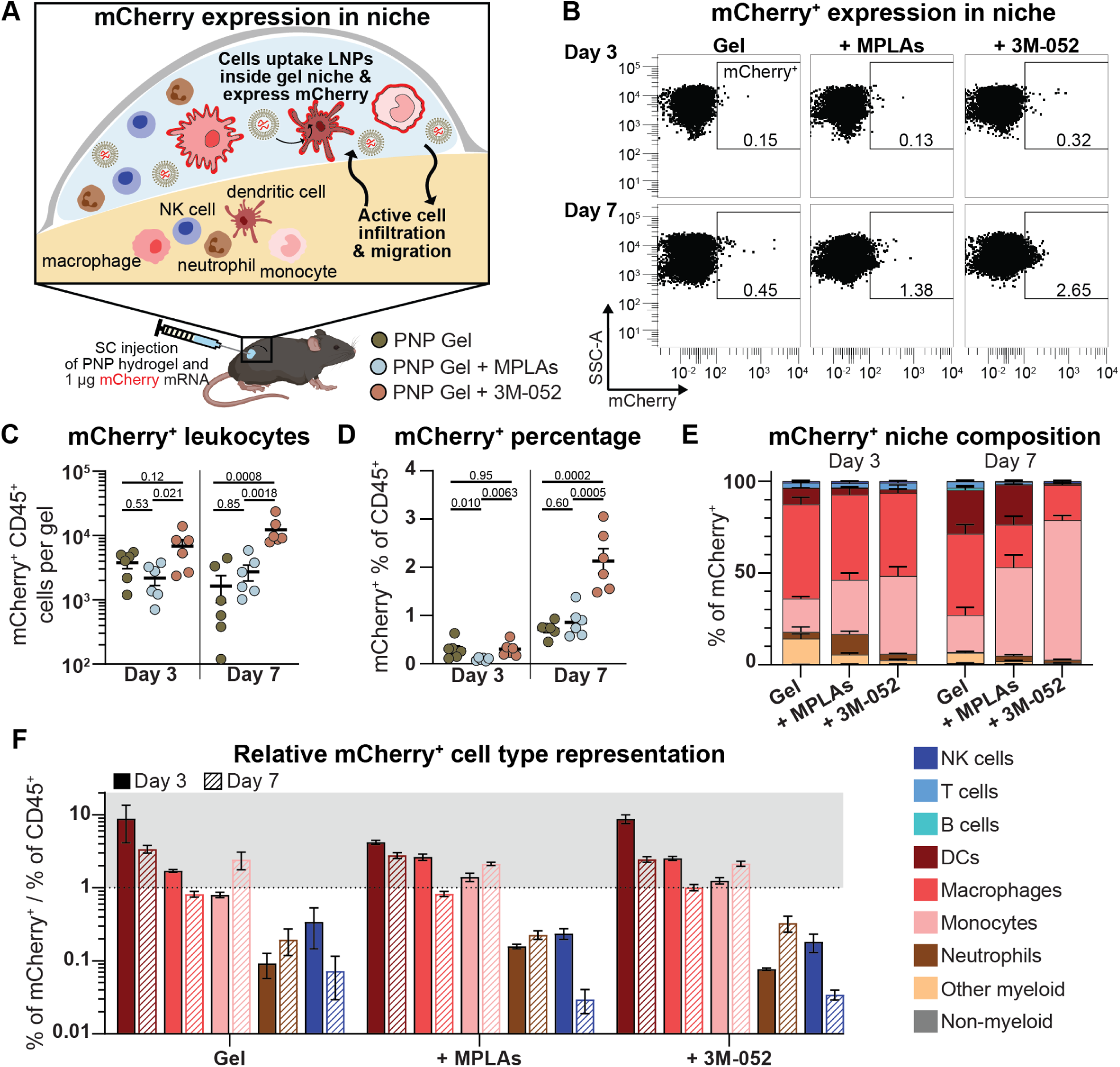
Characterization of mRNA expression in PNP hydrogel immune niche. A) Immune cells infiltrate injected PNP hydrogels loaded with mCherry mRNA/LNPs in vivo, take up LNPs, and express reporter mCherry protein. These cells can be quantified on days 3 and 7 following hydrogel excision with flow cytometry. B) Representative flow plots showing mCherry signal in CD45^+^ cells. C) The count of mCherry expressing CD45^+^ leukocytes per gel and D) those cells as a percentage of all CD45^+^ cells. Adjuvants increase the absolute number and percentage of CD45^+^ cells expressing the delivered mRNA. Data shown as mean ± SEM, n = 6. Statistical values shown are *p* values obtained from GLM fitting and Tukey HSD multiple comparison test in JMP. E) Quantification of cell types in the mCherry^+^ subniche per hydrogel as percentages of all gated CD45^+^ mCherry^+^ cells. Data shown as mean + SEM, n = 6. F) Ratio of the proportion each cell type makes up of the mCherry+ subniche to its percentage of the full CD45^+^ niche. Values of one indicate a cell is represented equally in the mCherry+ subniche as in the full gel. Values under one indicate fewer cells are mCherry^+^ than expected based on that cell type’s percentage of the CD45^+^ niche, and vice versa for values over one (overrepresented in mCherry^+^ subniche). Data shown as mean ± SEM, n = 6.

Considering the obvious differences observed in overall CD45^+^ cell niche compositions, yet relatively similar mCherry^+^ subniche compositions across groups, we sought to examine if certain cell types were specifically enriched in the mCherry^+^ subniche. To do this, we took the ratio of each cell type percentage of mCherry^+^ cells divided by the same cell type’s percentage of all CD45^+^ cells (**Figure 3F**). In this way, cell types which compose the same percentage of mCherry^+^ cells as they do all CD45^+^ cells would have a ratio of one. Similarly, ratios less than one would indicate fewer of that cell type are mCherry^+^ than expected given their frequency in the overall CD45^+^ niche, and vice versa for values over one (cell types enriched for mRNA expression). From this we observed APCs were overrepresented in the mCherry^+^ subniche while neutrophils and NK cells, despite composing a large fraction of the overall CD45^+^ cell niche, were underrepresented in the mCherry^+^ subniche. Looking at the fraction of each cell type that was expressing mCherry, less than 0.07% of NK cells and 0.76% of neutrophils expressed mCherry across all groups while up to 2.1, 4.5, and 5.3% of macrophages, monocytes, and DCs respectively expressed mCherry on day seven (**Supp Fig. S7**). These observations indicated that mRNA/LNPs incorporated into PNP hydrogels successfully transfected APCs, a key cell type for training adaptive immune responses. Further, the transfected cell types did not change substantially with different adjuvant cargo, whereas the surrounding cell niche did, providing a means to independently tune the context in which APCs express and present antigen. This approach presents a key opportunity when considering mounting different types of immune responses, such as anti-cancer or tolerogenic, where the context of antigen processing and presentation is critical.

### 2.4 Improved humoral and cellular response to commercial mRNA vaccine with PNP hydrogels

We next wanted to evaluate the impact of PNP hydrogels with and without adjuvants (and their corresponding cell niches) on the response to an mRNA/LNP vaccine. The modularity of PNP hydrogels allows simple incorporation of commercially available vaccines, like Moderna’s Spikevax vaccine encoding the full spike protein of SARS-CoV-2, in a simple off-the-shelf manner with and without adjuvants. We first evaluated two material formulations, PNP-0.5-5 and PNP-1-5, to assess any effects of material mechanical properties on the humoral response (**Supp Fig. S8**). PNP-1-5 hydrogels were an order of magnitude stiffer than PNP-0.5-5 hydrogels, resulting in a more persistent depot with slower release kinetics and potentially limited cellular infiltration at early time points. As mRNA degrades rapidly by hydrolysis, the softer materials performed better in vitro and in vivo, so we moved forward with the previously characterized PNP-0.5-5 formulation.

We delivered 0.25 μg mRNA/LNPs (Moderna Spikevax bivalent vaccine encoding WH1 and BA.4/.5) in a 100 μL subcutaneous injection in either PBS, the current clinical standard, or PNP-0.5-5 hydrogel with or without MPLAs or 3M-052. We dosed mice at weeks zero (prime) and eight (homologous boost) and collected serum for antibody titer evaluation over six months (**Figure 4A**). We immunized a separate cohort of animals and evaluated the T cell response via IFN-γ ELISpot at week ten (two-weeks post-boost). Serum titer ELISAs against WH1 spike protein at week eight post-prime showed that 5 in 6 animals exhibited measurable titers in the PNP hydrogel group compared to only 1 in 6 in the bolus control group (**Figure 4B**). Additionally, PNP hydrogel comprising 3M-052 produced significantly higher titers compared to both bolus and PNP hydrogel only (*p=*0.043 and 0.093 respectively). As early as week two, 3M-052 adjuvanted PNP hydrogel exhibited 4.5-fold improved antibody titers (*p*=0.029) while 3M-052 adjuvanted bolus had no observable benefit compared to unadjuvanted bolus (*p*=0.59) (**Supp Fig. S9**). At four months post-boost, animals in adjuvanted groups exhibited higher titers compared to bolus controls, and animals that received PNP hydrogel with 3M-052 exhibited titers nearly an order of magnitude higher than those that received bolus control or LNP-only hydrogel vaccines (*p* = 0.17 and 0.099 respectively) (**Figure 4C**). Antibody titers throughout the study showed that mRNA/LNP vaccine delivery in PNP hydrogels improved titers, particularly post-prime, and that PNP hydrogels adjuvanted with 3M-052 enhanced titers at all time points and produced substantively higher overall antibody exposure as measured by area under the curve (AUC) compared to the bolus control (*p*=0.071) (**Figure 4D, E**).

**Fig. 4.**
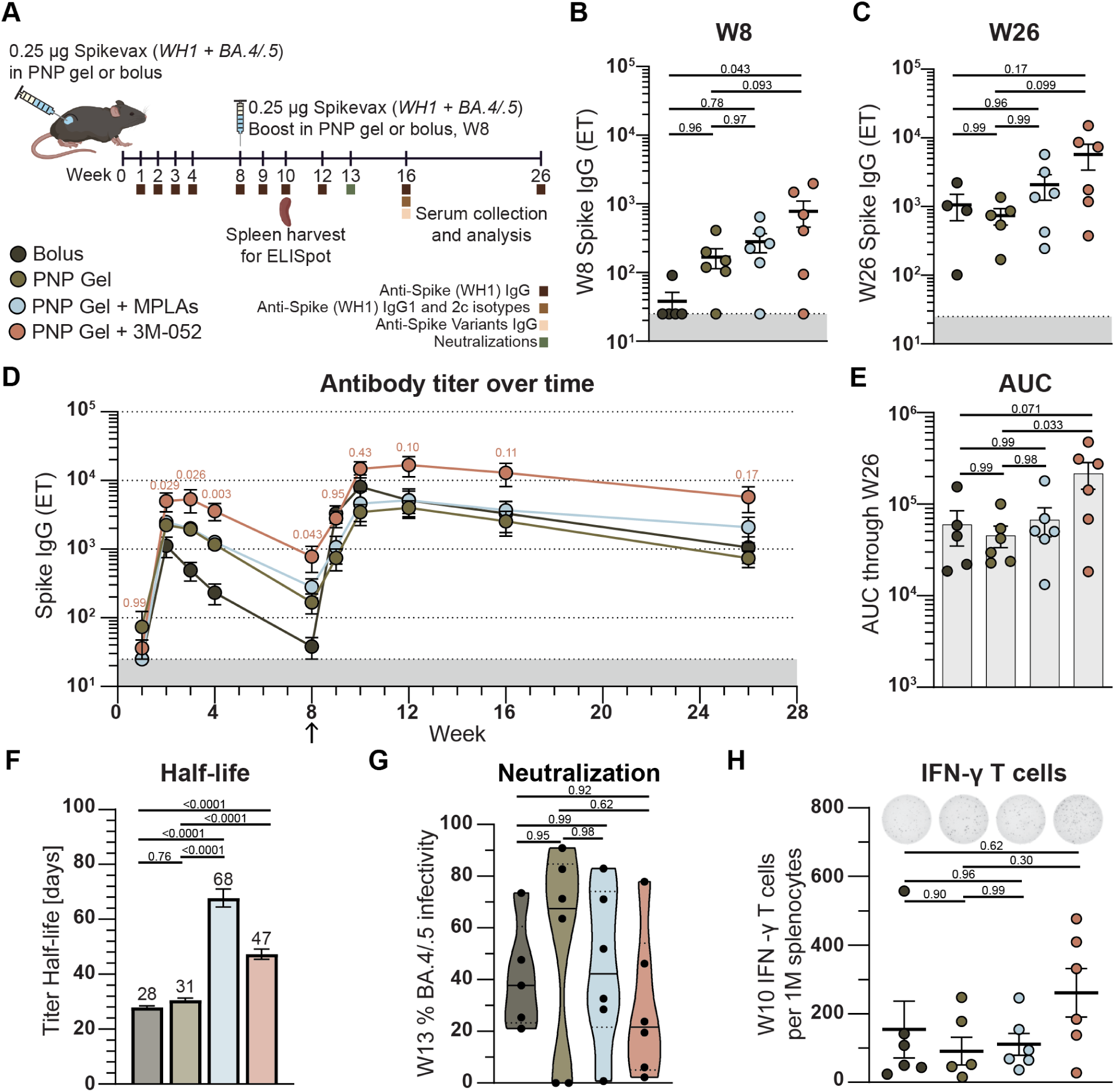
Humoral response to hydrogel-based mRNA/LNP vaccines. A) Mice were immunized with 0.25 μg commercially available bivalent SARS-CoV-2 mRNA vaccine (0.125 μg each variant) in PBS bolus or PNP-0.5-5 hydrogel with or without 1 μg 3M-052 or 20 μg MPLAs at week 0 and boosted with a homologous boost at week eight. Serum collected at designated time points was analyzed for anti-spike (Wuhan-Hu-1) antibodies via ELISA, as well as for other viral variant anti-spike antibodies, IgG subtypes, and neutralizing capacity. A separate cohort of animals primed and boosted in the same fashion were euthanized two weeks post-boost and spleens harvested for ELISpot evaluation of spike specific (WH1) T cells. Anti-WH1 spike IgG endpoint titers for all animals at B) week eight, C) week 26, and D) over time. E) Area under the curve of antibody titer from week 0 to 26 per animal. F) Decay half-life of antibody titer post-boost derived from parametric bootstrapping of titers following each treatment group’s post-boost peak. Data shown as mean ± SEM, n = 1000 simulations. G) Percent infectivity of BA.4/.5 pseudotyped lentivirus at week 13 post-prime and 1:50 serum dilution. H) Spike specific (WH1) IFN-γ producing splenocytes, as a proxy for T cells, at week two post-boost. Data shown as mean ± SEM, n = 4 - 6. Statistical values shown are *p* values obtained from GLM fitting and Tukey HSD multiple comparison test in JMP. In D, comparisons shown are to bolus control.

Considering durability is a concern with current commercial mRNA/LNP vaccines, we extracted an average antibody half-life using parametric bootstrapping on titer data following the post-boost peak (week 10 for bolus and week 12 for hydrogel groups, n=1000 runs) (**Figure 4F**). The PNP hydrogels increased antibody decay half-life by 11% over the bolus control, and adjuvanted PNP hydrogel groups increased antibody decay half-lives by 143% and 68% (MPLAs and 3M-052 respectfully, *p*<0.0001) over the bolus control. These enhancements represent a shift in half-life from one month for bolus delivery to upwards of 2.5 months for adjuvanted PNP hydrogels.

We next examined the functionality of the antibody response as well as the T cell response. We performed SARS-CoV-2 BA.4/.5 pseudotyped lentivirus neutralization assays at week thirteen post-prime (week five post-boost) and found no significant difference in infectivity between bolus or LNP-only hydrogels, and a trend toward enhanced neutralization (i.e., lower infectivity) in adjuvanted PNP hydrogel groups (**Figure 4G, Supp Fig. S10**). PNP hydrogels with 3M-052 exhibited a mean infectivity of 29.3% compared to 41.4 – 51.4% for other groups. Furthermore, only PNP hydrogel groups yielded infectivities below 10% at this dilution (2, 1, and 2 animals out of 6 treated animals for PNP only, with MPLAs, and with 3M-052, respectively). We next assessed functional T cell responses to WH1 spike peptides via ELISpot for IFN-γ producing splenocytes, a proxy for IFN-γ producing T cells, at week ten post-prime (week two post-boost). Vaccine delivery in PNP hydrogels alone was again comparable to bolus while adjuvanting with 3M-052 produced a more robust T cell response (**Figure 4H**). Altogether, the humoral and cellular responses to SARS-CoV-2 mRNA/LNP vaccines are improved with delivery in PNP hydrogels, and especially so with adjuvanted PNP hydrogels, whereas no improvement is achieved with standard bolus adjuvanting.

### 2.5 Increased breadth of humoral response to mRNA vaccine with PNP hydrogel delivery

Another important aspect of a vaccine response, particularly against highly mutable viruses like SARS-CoV-2, is the breadth of humoral responses. We evaluated antibody isotypes and endpoint titers at week sixteen post-prime against seven SARS-CoV-2 variants of concern, as well as SARS-CoV-1 (**Figure 5A, B**). Interestingly, all groups exhibited similar IgG1 titers, with PNP hydrogel groups showing modestly higher IgG1 titers compared to bolus, but only PNP hydrogels with 3M-052 produced measurable IgG2c titers in a majority of animals (4 in 6 treated animals). The IgG2c titer for PNP hydrogels with 3M-052 was significantly higher than that of PNP hydrogel only, as well as the bolus control (*p=*0.037 and 0.12, respectively). This observation was further reflected in the IgG2c/IgG1 ratio, a proxy for Th1 and Th2 skewing (**Figure 5C**). While PNP hydrogels exhibited comparable Th2 skewing to the bolus control, incorporation of adjuvants like 3M-052 elicited a clear skewing toward Th1. Since protection against different infectious diseases can be better mediated by different Th1/2 skewing depending on the disease, the ability to tailor the response in a simple and modular way by just changing the admixed adjuvant co-encapsulated within the PNP hydrogel is a significant benefit (*22*).

**Fig. 5.**
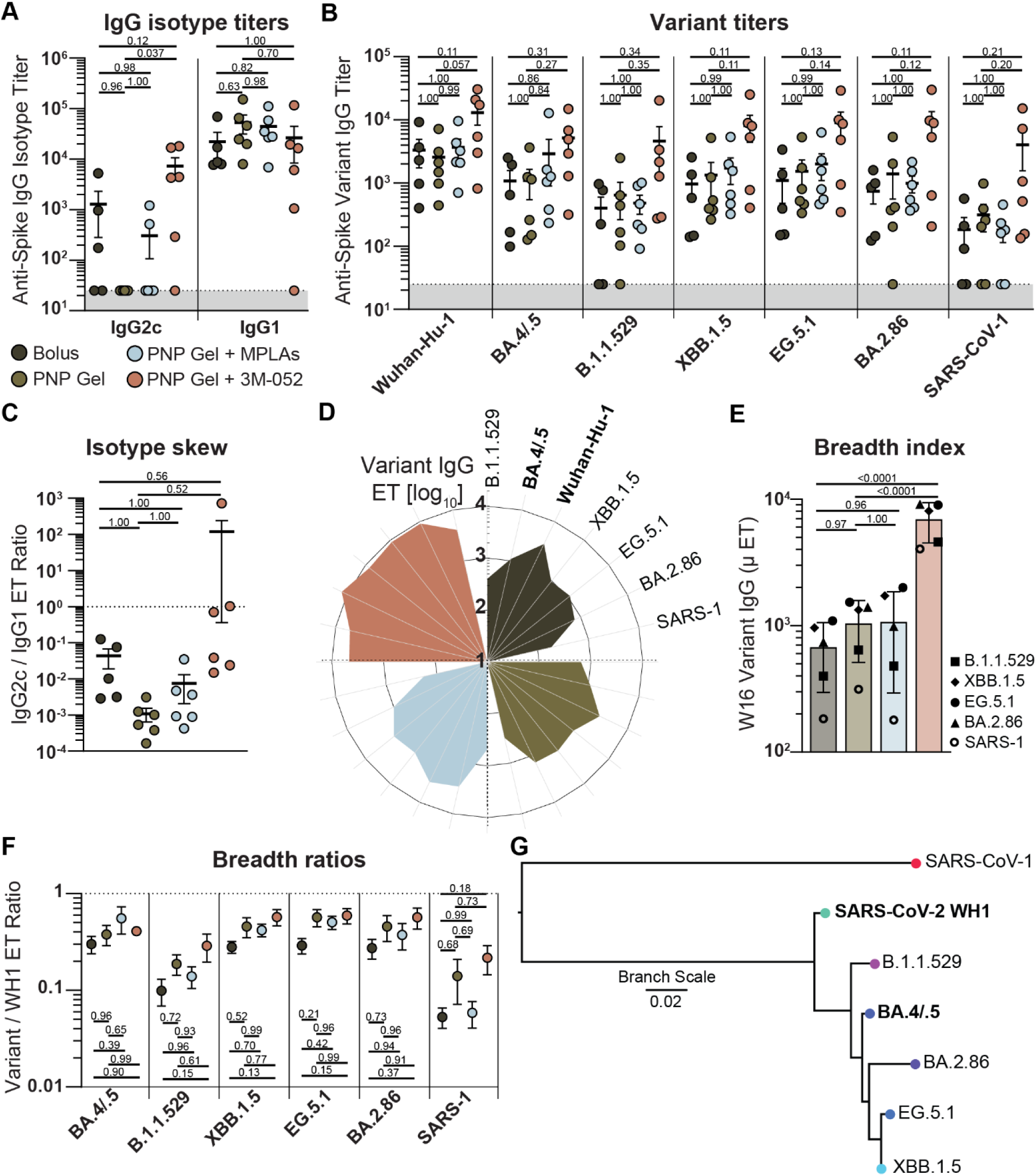
Breadth and functionality of response to hydrogel-based mRNA/LNP vaccines. A) IgG isotype endpoint titers at week 16 post-prime B) Anti-spike endpoint ELISA titers against different variants at week 16 post-prime. C) Ratio of IgG2c to IgG1 antibody isotypes indicating Th1 (higher values) or Th2 (lower values) skewing. D) Radar plot of absolute IgG endpoint titer for each variant. Broader and larger petals indicate improved breadth and consistency of response. E) Quantification of D, where each data point is the average of n = 5 - 6 animals against a single spike variant. Data shown as mean ± SD. F) Ratio of endpoint IgG titer for each variant compared with WH1 as a metric of consistency and breadth of humoral response. G) Phylogenetic tree showing lineage of variants assessed created using ‘Gene/Protein Tree’ tool on the Bacterial and Viral Bioinformatics Resource Center (BV-BRC) website https://www.bv-brc.org/. Unless otherwise written, data shown as mean ± SEM, n = 5 - 6. Statistical values shown are *p* values obtained from GLM fitting and Tukey HSD multiple comparison test in JMP.

Additionally, PNP hydrogel delivery of mRNA/LNP vaccines improved the breadth of antibody titers, particularly in adjuvanted PNP hydrogel groups. A petal/radar plot of anti-spike titers across variants is one way to quickly visualize increased breadth (**Figure 5D**). The PNP hydrogel group exhibited a slightly larger and broader petal than the bolus control, indicating an overall increase in antibodies against the tested variants not included in the vaccine, as well as a more consistent response across those variants. PNP hydrogel comprising 3M-052 elicited an even larger, broader petal than PNP hydrogel alone or with MPLAs. Of note, PNP hydrogels also reduced the number of non-responders to some variants like B.1.1.529 and SARS-CoV-1 over standard bolus vaccination, and PNP hydrogels with 3M-052 produced measurable anti-SARS-CoV-1 titers in all six animals. We further quantified the breadth of the humoral immune response with an absolute breadth index, composed of the average titer for animals in one treatment group against a single variant those animals were not vaccinated against (**Figure 5E**). This breadth index is slightly increased for PNP hydrogels alone and comprising MPLAs compared to bolus, and significantly higher for PNP hydrogels with 3M-052 (*p*<0.0001). We also evaluated the quality of breadth using a relative ratio comparing antibody titer for each variant to the titer against wildtype WH1 (**Figure 5F**). This breadth ratio analysis revealed that PNP hydrogel groups elicited more consistent responses, with ratios closer to one for all variants, indicating not just higher absolute titers but anti-variant titers closer to those against wildtype WH1 despite not being represented in the vaccine. The variants tested span CDC variants of concern and are genetically distinct within the Omicron clade (**Figure 5G**).

### 2.6 mRNA vaccination with PNP hydrogels improved magnitude and durability of germinal center response

To better understand the origin of the observed improvements in durability, breadth, and T cell responses imparted by PNP hydrogel delivery of mRNA/LNP vaccines with and without adjuvants, we used flow cytometry to probe the germinal center responses in draining lymph nodes at weeks one, two, and four following prime vaccination with 0.25 μg Moderna Spikevax bivalent vaccine (WH1 and BA.4/.5) (**Figure 6A, Supp Fig. S11**). Within the first week we observed that PNP hydrogel delivery elicited a higher proportion of activated B cells (MHCII^+^ CD86^+^), which was further increased with adjuvants and significantly so with 3M-052 (*p*=0.0003) (**Figure 6B**). We also found a significantly higher light zone to dark zone (LZ:DZ) ratio after one week for PNP hydrogel alone (*p*=0.09) and adjuvanted with 3M-052 (*p*=0.0003) compared to bolus vaccination (**Figure 6C**). These improvements observed only a week after prime injection in PNP hydrogel groups were interesting considering the slow release of cargo would lead us to expect slower germinal center kinetics compared to bolus. We hypothesize these early increases in lymph node and germinal center reactions result from the unique advantage of the in vivo immune niche formed within the PNP hydrogel depots and an early boost in the response from APCs transfected in the PNP niche.

**Fig. 6.**
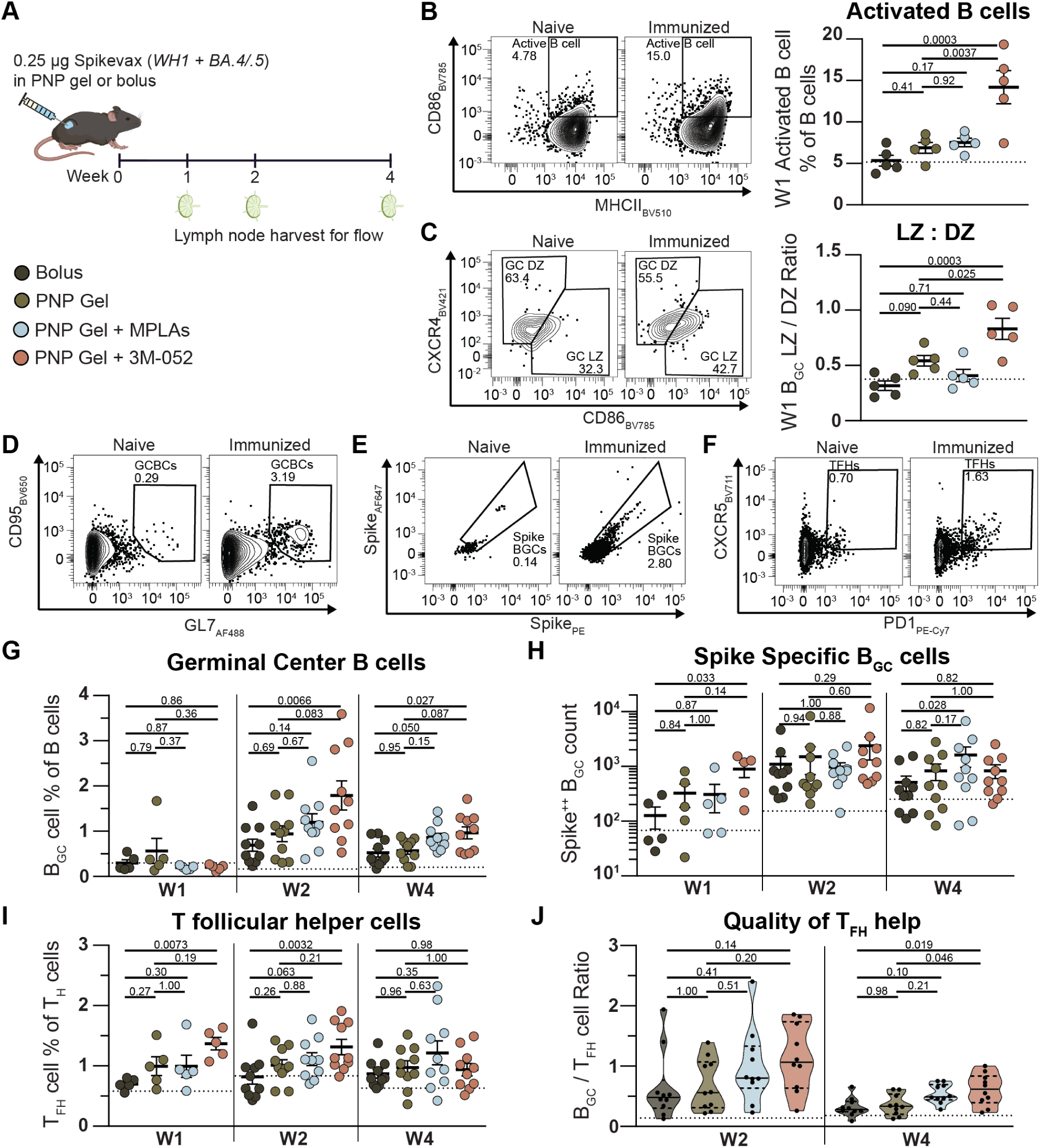
Germinal center response to hydrogel-based mRNA/LNP vaccines. A) Mice were immunized with bivalent SARS-CoV-2 mRNA vaccine in PBS bolus or PNP-0.5-5 hydrogel with or without adjuvants and lymph nodes harvested at weeks 1, 2, and 4. Week one representative flow plots and quantification of B) activated B cells (B220^+^ MHCII^+^ CD86^+^) and C) light and dark zone B_GC_ cells (B220^+^ CD95^-^ GL7^+^ CD38^-^ CD86^+^ CXCR4^-^ and CD86^-^ CXCR4^+^). D) Representative flow plots of B_GC_ cells (B220^+^ CD95^-^ GL7^+^), E) spike-specific B_GC_ cells (CD38^-^ Spike^++^), and F) T follicular helper cells (T_FH_, CD19^-^ CD3^+^ CD4^+^ CXCR5^+^ PD-1^+^). Percentages of parent population except Spike^++^ B_GC_ cells, which are of grandparent (see supplemental gating strategy). G) Quantification of B_GC_ cell percentage of all B cells. H) Counts of Spike^++^ B_GC_ cells. I) Quantification of T_FH_ cell percentage of all CD4^+^ T cells. J) Ratio of B_GC_ cells to T_FH_ cells as a metric of T_FH_ cell help quality. All data shown as mean ± SEM, n = 5 - 10. Statistical values shown are *p* values obtained from GLM fitting and Tukey HSD multiple comparison test in JMP.

We could more specifically examine the germinal center reaction by quantifying germinal center B (B_GC_) cells, including WH1 spike-specific B_GC_ cells, and T follicular helper (T_FH_) cells (**Figure 6D-F**). PNP hydrogels showed comparable or increased percentages of B_GC_ cells to bolus controls at all time points, with 0.29 – 0.69% in bolus and 0.56 – 0.94% in PNP hydrogel animals (**Figure 6G, Supp Fig. S12**). Adjuvanting with MPLAs provided a substantive increase over bolus or delivery in PNP hydrogels alone, with B_GC_ cells comprising up to 1.19% of B cells (*p*=0.14 and 0.67 at week two, *p*=0.050 and 0.15 at week four). PNP hydrogel adjuvanted with 3M-052 exhibited further significant improvements, up to 1.79% of B cells at week two, compared to bolus (*p*=0.0066 and 0.027 at weeks two and four), as well as a substantive benefit over unadjuvanted PNP hydrogel (*p*=0.083 and 0.087). We integrated B_GC_ cell percentages over time and observed PNP hydrogels increased the total B_GC_ cell exposure (AUC) by 36%, and PNP hydrogel groups adjuvanted with MPLAs and 3M-052 further increased this to 44% and 149% (*p*=0.19 for 3M-052) compared to bolus. We delved further into the B_GC_ cells and quantified those that were specific for the mRNA encoded antigen, WH1 SARS-CoV-2 spike protein. Spike-specific B_GC_ cells were indeed found to be increased in PNP hydrogel groups, significantly so in PNP hydrogels adjuvanted with 3M-052 (*p=*0.033 to bolus), and as early as week one, with persistence observed across the four weeks (**Figure 6H**). Interestingly, PNP hydrogels comprising MPLAs led to significantly more spike-specific B_GC_ cells at the later time point (week 4) compared to bolus (*p=*0.028), while the improvement in PNP hydrogel with 3M-052 was observed primarily in the earlier weeks.

Examining the T_FH_ cells, we observed an earlier response than B_GC_ cells, with higher percentages of T_FH_ cells at week one for PNP hydrogel (**Figure 6I, Supp Fig. S13**). T_FH_ cells comprised about 1% of CD4^+^ helper T cells for the PNP hydrogel group, but only 0.7 – 0.87% for the bolus group and upward of 1 – 1.22% and 0.94 – 1.37% for the PNP hydrogel groups adjuvanted with MPLAs and 3M-052. PNP hydrogels comprising 3M-052 maintained significantly higher percentages of T_FH_ cells than bolus at weeks one and two (*p*=0.0073 and 0.0032). PNP hydrogels comprising MPLAs also produced a substantively higher percentage of T_FH_ cells at week two than bolus (*p*=0.063), and this enhancement continued to increase through week four up to 1.37%, following a similar kinetic trend seen in the spike-specific B_GC_ cell counts across vaccine groups. Integration over time corroborated that PNP hydrogels increased T_FH_ cell percentage and overall exposure, and this was further increased for PNP hydrogels with adjuvants.

The ratio of B_GC_ cells to T_FH_ cells can be used as a metric for the quality of T_FH_ cell help, which we evaluated at weeks two and four (we excluded week one due to the weaker B_GC_ response) (**Figure 6J**) (*44*). More animals exhibited consistently higher quality T_FH_ cell help in the PNP hydrogel group at week two, with 4 in 10 animals at or above the third quartile compared with only 2 in 10 animals in the bolus group, which also had a lower third quartile value. Adjuvanting of the PNP hydrogels with MPLAs or 3M-052 further improved the quality of T_FH_ cell help, both in magnitude (average ratio) as well as number of animals experiencing better help (i.e., the spread of animals above the average). Week four ratios showed a slight waning, but adjuvanted PNP hydrogel delivery prolonged significantly better quality T_FH_ cell help compared to bolus (*p*=0.10 and 0.019 for MPLAs and 3M-052) and PNP hydrogel alone (*p*=0.21 and 0.046 for MPLAs and 3M-052). It was clear PNP hydrogels, particularly PNP hydrogels adjuvanted with 3M-052, improved the magnitude, duration, and quality of the germinal center reaction to mRNA/LNP vaccines, which reflected improvements in antibody titers, breadth, and durability found earlier.

## 3. Discussion

In this work we showed that delivering a commercially available SARS-CoV-2 mRNA/LNP vaccine within an injectable PNP hydrogel depot led to improved humoral and cellular immune responses driven by increased and prolonged germinal center reactions. PNP hydrogels enable facile formulation of off-the-shelf mRNA/LNPs with various adjuvant cargos, create tunable immunological niches in vivo with different adjuvant cargos, and promote transfection of APCs within the niche without any modifications to the LNPs. We characterized incorporation of mRNA/LNPs into PNP hydrogel materials and showed that formulation in this way does not impact material properties nor LNP transfection efficiency. We then characterized the cellular milieu within the PNP hydrogel depot in vivo and determined it to be highly plastic and tunable based on incorporation of adjuvant cargo. We further showed that key APCs within this niche preferentially expressed delivered mRNA. Following this characterization, we showed that commercial mRNA/LNP vaccine delivery in a PNP hydrogel niche with and without adjuvants improved the efficacy of Moderna Spikevax over clinically relevant bolus controls. Finally, we investigated the germinal center reactions driving these observed improvements and found that PNP hydrogel delivery increased the magnitude, duration, and quality of these reactions.

In contrast to chemical strategies to target LNPs to APCs, our approach does not require altering LNPs and can be readily applied to commercially available mRNA/LNP vaccines. Furthermore, these PNP hydrogels may be more widely applicable to other mRNA and nucleic acid delivery platforms like poly(β-amino esters) (PBAEs) or viral vectors, although this would require further study (*3*). Interestingly, we found the best humoral response and most active germinal center reactions in animals immunized with PNP hydrogels adjuvanted with 3M-052, a potent TLR7/8 agonist, yet this hydrogel niche was found to host the fewest DCs and instead comprised substantially more transfected monocytes than other tested groups (**Fig. 3, 4**). We initially hypothesized better vaccine responses would positively correlate with the presence of DCs in the PNP hydrogel niche; however, these data indicate this is not necessarily the case and highlight an important gap in knowledge in the field. Indeed, there is little data characterizing cellular recruitment to the injection site or understanding correlations between injection site cellular milieu and strong or specific mounted immune responses. This shortfall is due in part to fundamental difficulties in measuring cells within the injection site, particularly in the subcutaneous space. While some research has attempted to characterize the immune infiltrate following intramuscular or intradermal injections, these studies have relied heavily on imaging modalities and immunohistochemistry approaches requiring biopsies of an injection site which can be difficult to find reproducibly (*8, 45*). One of the key advantages of the PNP platform is that it allows us to study injection site reactions with single cell resolution in a way previously not possible. Future work could consider additional processing, like single cell sequencing and cytokine profiling, to further elucidate differences at the injection site which may correlate to overall shifts in the downstream immune responses.

While adjuvants are known to improve the immune response to protein antigens, adjuvanting mRNA vaccines is complicated and under-characterized, requiring a balanced approach as excessive immune activation and cytokine production can hamper mRNA translation (*46*). Adjuvants, particularly TLR agonists, exhibit a wide range of physiochemical properties, from small molecules to large single or double stranded nucleic acid chains, which make their interactions with mRNA vaccines difficult to study in a systematic or modular fashion. It is important to more thoroughly understand the impact of adjuvants on mRNA vaccines, as well as to find adjuvants which improve and tailor vaccine responses for different antigens or diseases. Prior works adjuvanting mRNA vaccines have focused on incorporating adjuvants into LNPs, often requiring modifications to both the adjuvant and the particle formulation and are generally nontransferable to other adjuvant molecules. The PNP hydrogel platform provides a unique means to independently incorporate adjuvants alongside mRNA/LNPs, allowing for unprecedented control over adjuvanting mRNA vaccines and investigating the impact on immune responses. In this work, we find that TLR4 activation by MPLAs has little benefit over PNP hydrogel delivery alone, but TLR7/8 activation by 3M-052 significantly improves the magnitude and quality of vaccine responses. Interestingly, PNP hydrogels alone are rather inert and do not modulate T-helper skewing (**Fig. 5**) like other depot technologies such as Alum, which produces a strong Th2-skewed response even when combined with Th1-skewing adjuvants like 3M-052 (*47*).

Future work could explore other adjuvants targeting different TLRs, such as TLR1/2a (Pam3CSK4), TLR9a (CpG), or STING (cGAMP), as well as combinations of adjuvants shown in other contexts to afford synergistic immune activation (*22, 48, 49*). Additionally, the PNP hydrogel platform enables encapsulation of other immune activating cargo in a plug-and-play manner, like protein chemokines or cytokines, that would otherwise be difficult or impossible to co-deliver with mRNA/LNPs without modification (*33*). Granulocyte-macrophage colony stimulating factor (GM-CSF) is a small protein which should drive infiltration of more DCs into the PNP material and could be combined with other adjuvants. Cytokines like IL-10 or IL-2 could also be incorporated to induce a more tolerogenic niche in the PNP hydrogel (*50, 51*). The modularity of this platform can effectively skew the immunological niche and subsequent response based on encapsulated molecular cargos, creating opportunities beyond improving infectious disease vaccines to directing improved cytotoxic responses in cancer vaccines or suppressive responses in tolerogenic applications.

While PNP delivery of mRNA/LNPs offers a lot of potential to improve vaccine efficacy as well as introduce a platform to study the impact of adjuvants and the injection site kinetics in a way otherwise unstudied, there remains work to be done. One limitation of the presented work is that it focuses on a single target disease, SARS-CoV-2, and does not expand to other antigens that may show different responses. Additionally, the PNP hydrogel niche is characterized at two time points, but mRNA expression is transient and some immune cells have very short lifespans (e.g., only 3 days for neutrophils) (*52*). Follow up work should evaluate the kinetics of the immune niche and mRNA expression in greater depth and at additional time points, as well as investigate the kinetics of antigen and cellular accumulation in the lymph nodes. Furthermore, this work could be conducted in Ai14/Cre mice to allow analysis of all cells which translate delivered mRNA, not just the ones currently expressing enough protein to be detected (*53*). This would further allow decoupling of cellular expression of encoded protein from the possibility that cells within the PNP niche are phagocytosing protein made by other transfected cells in the niche.

Taken together, the presented work shows that PNP hydrogel delivery of mRNA/LNPs is a promising strategy to employ a tunable immunomodulatory platform for a variety of applications, ranging from probing fundamental questions about injection site cellular reactions to mRNA vaccines, to improving vaccine efficacy against infectious diseases without altering LNP chemistries, and even potential to tailor pro-inflammatory or tolerogenic responses.

## 4. Experimental Methods

### PNP hydrogel formulation

HPMC-C_12_ and PEG-PLA was prepared as described in supplemental methods. HPMC-C_12_ was dissolved at 2 wt% in PBS and loaded into a 1 mL luer-lock syringe. A 20 wt% solution of PEG-PLA NPs in PBS was added to a solution of PBS with or without mRNA/LNPs and/or adjuvant depending on formulation, and loaded into a second 1 mL syringe. The two syringes were connected with a female-female luer lock elbow, with care to avoid air at the interface of the HPMC-C_12_ and nanoparticle solution, and gently mixed until a homogenous PNP hydrogel was formed. Hydrogels were formulated with final concentrations of 0.5 or 1 wt% HPMC-C_12_ and 5 wt% NPs, denoted PNP-0.5-5 or PNP-1-5.

### PNP hydrogel rheological characterization

Rheological characterization was performed on PNP hydrogels with or without Moderna Spikevax monovalent (50 μg mL^-1^) using a TA Instruments DHR-2 stress-controlled rheometer. All experiments were performed using a 20 mm diameter serrated plate geometry at 25 °C with a 500 µm gap. Frequency sweep measurements were performed at a constant 1% strain in the linear viscoelastic regime. Stress sweeps were performed from low to high with steady state sensing and yield stress values defined as the stress at the intersection of lines tangent to the plateau and the yielding regimes. Flow sweeps were performed from high to low shear rates. Step shear experiments were performed by alternating between a low shear rate (0.1 s^−1^; 60 s) and a high shear rate (10 s^−1^; 30 s) for three cycles.

### In vitro LNP transfection

RAW-Blue cells (InvivoGen, raw-sp) were cultured at 37 °C with 5% CO_2_ in DMEM suplemented with high glucose, L-glutamine, sodium pyruvate (Cytiva, SH30243.FS), 10% heat inactivated fetal bovine serum (Cytiva, SH30396.03HI), and penicillin (100 U mL^−1^)/streptomycin (100 μg mL^−1^)(Cytiva, SV30010). 100 k cells were plated in each well of a tissue culture treated 96-well plate in 200 μL media and allowed to adhere for 24 h. After 24 h, media was aspirated, replaced with 180 μL fresh media, and 20 μL of PNP hydrogel or PBS with 5 μg mL^−1^ luciferase mRNA/LNPs (collaborators in the Irvine group at MIT) was injected through a 26-gauge needle into each well (n=3-4 wells) for a final concentration of 0.1 μg per well. The plate was incubated for 24 hours and transfection efficiency measured using Bright-Glo^TM^ Luciferase Assay System (Promega). Briefly, 100 μL of BrightGlo reagant was added to each well and, after a 3 minute incubation, 180 μL from each well was transferred to a white 96 well plate and luminescence was read on a Synergy H1 Microplate Reader (BioTek Instruments).

### Mice and vaccination

All animal studies were performed in accordance with the National Institutes of Health (NIH) guidelines, with the approval of the Stanford Administrative Panel on Laboratory Animal Care. Seven to eight week old female C57BL/6 were purchased from Charles River and housed in the animal facility at Stanford University. Mice were shaved and injected through a 26-gauge needle subcutaneously on the right flank with 100 μL of either bolus or gel vaccine under brief isoflurane anesthesia. Unless noted otherwise, mice were injected within five weeks of arriving at Stanford. Blood was collected weekly from tail veins. For flow cytometry, mice were euthanized under CO_2_ and organs such as inguinal lymph node (LN) and spleen were explanted or hydrogels were explanted from the subcutaneous space.

### Vaccine and other formulations

Vaccine primes and boosts (eight weeks after prime) contained 0.25 μg Moderna Spikevax bivalent (WH1 and BA.4/.5, 0.125 μg each variant) per dose in either bolus (PBS) or in PNP hydrogels with or without 20 μg per dose MPLAs (Invivogen, vac-mpls) or 1 μg per dose 3M-052 (AAHI). Spikevax was obtained from the Stanford Hospital Pharmacy and stored at 4 °C until use within seven days of first septa puncture, or frozen at −20 °C and only thawed once before use. For gel infiltration studies, 1 μg per dose of mCherry encoding mRNA/LNPs was incorporated into PNP hydrogels and dosed. mRNA/LNPs encoding mCherry or firefly luciferase were prepared via nanoprecipiation and commercially available lipids mimicking the Moderna Spikevax formulation. Briefly, SM-102 (BroadPharm), DSPC (Avanti Polar Lipids), cholesterol (Avanti Polar Lipids), and PEG-DMG 2000 (Avanti Polar Lipids) were dissolved in ethanol in the molar ratio 50:10:38.5:1.5 to make a lipid solution. mRNA with 5-methoxyuridine (5moU) modification encoding mCherry or firefly luciferase (TriLink) was dissolved in citrate butter (pH 3). LNPs were formulationed by mixing lipid and mRNA solutions in the ratio 1:3 (v/v) at 12 mL min^-1^ using an Ignite NanoAssemblr (Precision Nanosystems). LNPs were dialized overnight against 20 mM Tris acetate and 8% (w/v) RNAse-free sucrose (VWR) using 3,500 MWCO dialysis cassets (Thermo Fisher), aliquoted, and stored at −80 °C until use.

### Flow cytometry of PNP hydrogel immune infiltrate

Mice were shaved and injected subcutaneously on the right flank with 100 μL PNP-0.5-5 hydrogels with 1 μg dose^−1^ mCherry mRNA/LNPs, with or without MPLAs or 3M-052. At three and seven days after injection, mice were euthanized by CO_2_ and hydrogel depots extracted and placed in microcentrifuge tubes with 750 μL FACS buffer (PBS, 3% heat inactivated FBS, 1 mM EDTA). Hydrogels were mechanically disrupted to single cell suspensions using Kimble BioMasherIIs (DWK Life Sciences). Suspensions were passed through a 70 μm cell filter (Celltreat, 229484) into 15 mL Falcon tubes, spun at 500 rcf for 5 minutes, resuspended in PBS, and counted using acridine orange/propidium iodide cell viability stain (Vitascientific, LGBD10012) and a Luna-FL dual Fluorescence cell counter (Logos biosystems). 1 million live cells per sample were transferred to a 96-well conincal bottom plate (Thermo Scientific, 249570) and stained.

### Mouse serum ELISAs

Antigen-specific IgG endpoint titers were measured using an endpoint ELISA. MaxiSorp plates (Thermo Scientific, 449824) were coated with SARS-CoV-2 Spike trimer (Sino Biological, 40589-V08H4) or variant trimer (Sino Biological, 40589-V08H32, 40589-V08H26, 40589-V08H45, 40589-V08H55, 40589-V08H58, or 40634-V08B) at 2 μg mL^-1^ in PBS at 4 °C overnight and subsequently stored at −80 °C. Plates were thawed for 1-2 h at room temperature, washed 5 times with 300 μL per well wash solution (PBS with 0.05% Tween 20), and blocked with diluent buffer (PBS with 1% bovine serum albumin) for 2 h. All incubation steps were at room temperature on a rotator and plates were washed 5 times with wash solution between each step. Serum dilutions were prepared during the blocking step in a conical bottom plate (Thermo Scientific, 249570) in diluent buffer starting at 1:100 (1 μL serum into 99 μL diluent buffer) and serially diluted 4-fold. Following blocking and washing, serum dilutions were transferred to antigen-coated plate, 50 μL per well, and incubated for 2 hr. Goat anti-mouse IgG Fc HRP (Invitrogen, A16084) was diluted from glycerol stock (1 mg mL^-1^) into diluent buffer (1:10,000) and 50 μL added to each well and incubated for 1 h. For isotype ELISAs, goat anti-mouse IgG1 or IgG2c heavy chain HRP (Abcam, ab97240 or ab97255) were diluted 1:20,000 from vendor stock solution. Plates were developed for 6 minutes with high sensitivity TMB substrate (Abcam, ab171523) and reaction was stopped with 1 N HCl. Absorbance was read at 450 nm with a Synergy H1 microplate reader (BioTek Instruments). Data were analyzed in GraphPad Prism and fit with a five point asymmetric sigmoidal curve with constraints S > 0, top < 4, and bottom = 0.048 (background absorbance value). The endpoint titer was defined as the serum dilution value at which the absorbance reached 2x the background value, or 0.1, interpolated from the curve fits in GraphPad Prism. For isotype plates, background absorbance was set to 0.094 and 0.066 for IgG1 and 2c respectively and endpoint was defined at 0.2. Samples below endpoint absorbance at 1:100 dilution were designated below the limit of detection and the endpoint titer defined as a dilution value of 25.

### Serum neutralization assays against pseudoviruses

Antisera were heat inactivated (56 °C, 30– 60 min) before neutralization assays. Neutralization against SARS-CoV-2 BA.4/5 was analyzed in HeLA-ACE2/TMPRSS2 cells. One day before infection (day 0), cells were seeded at 8,000 cells per well in white-walled, white-bottom, 96-well plates (Thermo Fisher or Greiner Bio-One). On day 1, antisera were serially diluted in D10 media and mixed 1:1 with pseudoviruses for 1-2 hrs at 37 °C before being transferred to cells. The pseudovirus mixture contained SARS-CoV-2 BA.4/5, D10 media and polybrene (1:500). Assays were read out with luciferase substrates 2 d after infection by removing the media from the wells and adding 80 uL of a 1:1 dilution of BriteLite in DPBS (BriteLite Plus, Perkin Elmer). Luminescence values were measured using a microplate reader (BioTek Synergy™ HT or Tecan M200). Percent infection was normalized on each plate. Neutralization assays were performed in technical duplicates.

### ELISpot

Spike-specific IFN-γ producing splenocytes were evaluated using a Mouse IFN-γ Single color ELISpot Kit (CTL) and CTL counter (Immunospot S6 Ultra M2). Spleens were harvested from mice two weeks post-boost, disrupted to single cell suspensions with frosted glass slides in CTL test media (CTLT-005) supplemented with 1% L-glutamine (termed CTL-g), passed through 70 μm cell filters into 15 mL Falcon tubes, and spun at 400 rcf for 4 minutes. Red blood cells were lysed with 1 mL ACK lysis buffer (Gibco, Thermo Scientific, A1049201) for 2 minutes and reaction was quenched with 9 mL CTL-g. Samples were spun at 400 rcf for 4 minutes, resuspended in CTL-g, counted, and plated in a low-bind 96-well conincal bottom plate. Samples were transferred to pre-coated ELISpot plates and stimulated for 24 h at 37 °C with either 5 μg peptide^-1^ mL^-1^ SARS-CoV-2 pan-spike peptides (JPT Peptide Technologies, PM-WCPV-S-1), 1 μg mL^-1^ concanavalin A (ConA, Sigma-Aldrich, C0412-5MG) as a positive control, or CTL-g as a negative control. Cells were at a final concentration of 200 k in negative wells, 400 k in ConA wells, and both 200 k and 400 k in peptide wells. After 24 h, spots were developed following manufacturer’s instructions.

### Flow cytometry of lymph nodes

Draining inguinal lymph nodes were harvested from mice at weeks one, two, and four post-prime. LNs were disrupted, filtered, and counted as previosly described in hydrogel preparation and 1 million live cells per sample were transferred to a 96-well conincal bottom plate and stained.

### Staining protocol and panels

LN and hydrogel samples were first stained with 100 μL Live/Dead Fixable Near-IR (Thermo Scientific, L34975) for 30 minutes at room temperature, quenched with 100 μL FACS buffer, and spun at 935 rcf for 2 minutes. Samples were then incubated with 50 μL anti-mouse CD16/CD32 (1:50 dilution; BD, 553142) for 5 minutes on ice before incubating with 50 μL full antibody stain for 40 minutes on ice for hydrogel samples and 60 minutes for lymph node samples. Samples were spun and resuspended in 50 - 70 μL FACS buffer and run on the BD FACSymphony A5 SORP in the Stanford Shared FACS Facility. Data was analyzed in FlowJo.

LN full antibody stain included anti-CD184 (1:400 dilution; BV421; Fisher Scientific, 50-605-189), anti-CD138 (1:400; BV605; Fisher Scientific, 50-207-1399), anti-CD86 (1:400; BV785; Fisher Scientific, NC1188484), anti-CD279 (1:400; PE-Cy7; BioLegend, 135215), anti-CD19 (1:200; PerCP/Cy5.5; Fisher Scientific, 50-113-0313), anti-CD95 (1:200; BV650; Fisher Scientific, BDB740507), anti-CD38 (1:200; BUV737; BD, 741748), anti-CD4 (1:200; BUV805; BD, 612900), anti-I-A/I-E (1:200; BV510; BioLegend, 107636), anti-GL7 (1:100; AF488; Fisher Scientific, 50-711-897), anti-CD3 (1:100; AF700; Fisher Scientific, 50-162-375), anti-CD45R (1:100; BUV395; BD, 563793), anti-CD185 (1:50; BV711; Fisher Scientific, 50-207-1644), anti-spike tetramer (20nM; AF647), and anti-spike tetramer (20nM; PE). Tetramers were prepared on ice by adding AF647- or PE-Streptavidin (Thermo Scientific, S32357 or BD, 554061) to 6 μM biotinylated wildtype SARS-CoV-2 spike trimer (Sino Biological, 40589-V27B-B) in five steps, once every 20-60 min, for a final molar ratio of 4.1:1 spike protein to dye and concentration of 0.5 μM.

Hydrogel full antibody stain included anti-I-A/I-E (1:800 dilution; FITC; BioLegend, 107605), anti-CD45 (1:800; AF700; BioLegend, 103127), anti-Ly6C (1:400; BV570; BioLegend, 128029), anti-XCR1 (1:200; AF647; BioLegend, 148213), anti-F4/80 (1:200; BV421; BioLegend, 123137), anti-Ly6G (1:200; BV711; BioLegend, 127643), anti-CD3e (1:200; PerCP-eFluor710; Thermo Scientific, 46-0033-82), anti-CD19 (1:200; PE-Cy7; BioLegend, 115519), anti-CD11c (1:200; PE; BioLegend, 117307), anti-CD11b (1:100; BV510; Fisher Scientific, 50-112-9846), and anti-NK1.1 (1:100; BV605; BioLegend, 108753).

### Statistical Analysis

For in vivo experiments, animals were cage blocked in all experiments except the humoral study and data presented as mean ± SEM. Comparisons between multiple groups were conducted with the general linear model (GLM) and Tukey HSD test in JMP, accounting for cage blocking. In order to normalize the variance and meet requirements for statistical tests used, flow cytometry data presented as percentages (e.g. B_GC_ cells percent of B cells) were transformed using the equation y = ln(x/(100-x)), where x is the original data point plotted and y the data statistics are run on. Comparisons between two groups were conducted with multiple unpaired two-tailed student t-tests run in GraphPad Prism with false discovery rate (FDR) correction using two-stage step-up method of Benjamini, Krieger. Select *p* values are shown in the text and figures and all *p* values are in the supporting information.

## Acknowledgements

The authors would like to thank every member of the Appel Lab, former and current, for their on-going support, technical expertise, and scientific discussion. In particular, the authors thank Changxin (Lyla) Dong, Alakesh Singh, Priya Ganesh, Ibukun Ajifolokun, Jerry Yan, and Leslee T. Nguyen for their assistance with tissue dissociation. The authors also thank Dr. Volker Böhnert for help designing an initial flow cytometry panel for PNP hydrogel cell infiltration.

## Author contributions

Conceptualization: ELM, EAA

Methodology: ELM, AU, BSO

Software: NE, ELM

Investigation: ELM, JHK, CKJ, AU, JB, YES, OMS, SCW

Resources: TM, NC, DJI

Visualization: ELM

Supervision: EAA

Writing—original draft: ELM

Writing—review & editing: JHK, SCW, JB, TM, NC, DIJ, EAA

## Data and materials availability

All data needed to evaluate the conclusions of this paper are present in the main text and/or the supplementary materials.

