## Supplemental Information for "Generation of an inflammatory niche in an injectable hydrogel depot through recruitment of key immune cells improves efficacy of mRNA vaccines"

### 1. Supplementary Methods

*Materials:* Hypromellose (HPMC, meets USP testing specifications), N-methyl-2-pyrrolidone (NMP), 1-dodecylisocynate, N,N-diisopropylethylamine (DIPEA), acetone, monomethoxy-PEG (5 kDa), diazobicylcoundecene (DBU), acetic acid, diethyl ether, hexanes, dimethyl sulfoxide (DMSO), acetonitrile were purchased from Sigma-Aldrich and used as received. Dichloromethane (DCM) was purchased from Sigma-Aldrich and further dried via cryo-distillation. Lactide was purchased from Sigma-Aldrich and recrystallized from ethyl acetate (dried over sodium sulfate) three times.

*HPMC-$C_{12}$ synthesis:* HPMC−$C_{12}$ was prepared according to previously reported procedures (*34*). HPMC (1.0 g) (SEC MALS: Mw (Đ) = 372.4 kDa (1.43), method previously reported) was dissolved in NMP (40 mL) at room temperature with stirring. Once the polymer had completely dissolved, the reaction was brought to 50 °C and a solution of 1-dodecylisocynate (0.5 mmol) in NMP (5 mL) was added dropwise, followed by DIPEA (catalyst, 125 µL). The reaction was maintained at 50 °C for 30 minutes, then heat was shut off and mixture was left stirring overnight at room temp. The solution was then precipitated from acetone and HPMC-$C_{12}$ was purified by dialysis against MilliQ water for 3-4 days (MWCO 3.5 kDa) and lyophilized, yielding HPMC-$C_{12}$ as a white amorphous powder. The polymer was dissolved at 20 mg $mL^{-1}$ in sterile PBS, pH 7.4, prior to use in hydrogels.

*PEG-PLA synthesis:* PEG-PLA was prepared and analyzed as previously reported (*34*). Recrystallized lactide (10 g) was fully dissolved in cryo-distilled DCM (45 mL) under $N_{2\,(g)}$ with mild heating. Methoxy poly(ethylene glycol) (5 kDa; 2.5 g) was heated to 100 °C under vacuum for 1-2 h, allowed to cool under $N_2$, and then dissolved in cryodistilled DCM (5 mL). Once dissolved, the full PEG solution was added to the lactide solution under $N_2$ and mixed with hand swirling. A solution of DBU (150 µL cryodistilled DBU per 1 mL cryodistilled DCM) was prepared and 500 µL added to the lactide-PEG solution under $N_2$. The reaction was swirled by hand and allowed to react for 8 min before quenching with acetic acid (~2 drops in 500 µL acetone). The PEG-PLA copolymer was precipitated from excess 50:50 mixture ethyl ether and hexanes, collected, and dried under vacuum to yield a white amorphous powder. DMF GPC: Mw (Đ) = 24.5 kDa (1.13), method previously reported (*34*).

*PEG-PLA nanoparticle (NP) preparation*: NPs were prepared and analyzed as previously reported (*34*). Briefly, a solution (1 mL) of PEG-PLA in 25:75 DMSO:acetonitrile (50 mg $mL^{-1}$) was added dropwise to water (10 mL) at a stir rate of 600 rpm. NPs were purified by ultracentrifugation over a filter (MWCO 10 kDa; Millipore Amicon Ultra-15) followed by resuspension in PBS to a final concentration of 200 mg $ml^{-1}$. NP size and dispersity were characterized by DLS (Wyatt DynaPro PlateReader-II; average diameter = 34.1 nm, PDI = 0.05)

*Pseudotyped lentivirus production:* SARS-CoV-2 spike-pseudotyped lentiviruses encoding a luciferase-ZsGreen reporter were produced in HEK293F cells by co-transfection of five plasmids. The five plasmids include a packaging vector (pHAGE-Luc2-IRES-ZsGreen), a plasmid encoding the SARS-CoV-2 spike (HDM-SARS2-spike-delta21, Addgene, 155130) and three helper plasmids (pHDM-Hgpm2, pHDM-Tat1b and pRC-CMV_Rev1b). BA.4/.5 mutations include T19I, Δ24-26, A27S, G142D, V213G, G339D, S371F, S373P, S375F, T376A, D405N, R408S, K417N, N440K, L452R, S477N, T478K, E484A, F486V, Q493R, Q498R, N501Y, Y505H, D614G, H655Y, N679K, P681H, N764K, D796Y, Q954H, N969K. 50 mL of cells were diluted to a density of approximately 3–4 × $10^6$ cells per mL. Transfection mixture was prepared by adding five plasmids (50 µg of packaging vector, 17 µg of SARS-CoV-2-encoding plasmid and 11 µg of each helper plasmid) to 5 mL of Expi-free medium, followed by the dropwise addition of BioT transfection reagent (150 µl, Bioland Scientific) with vigorous mixing. After 10 min incubation at room temperature, the transfection mixture was transferred to HEK293F cells. D-glucose (4 g $L^{-1}$, Sigma-Aldrich) and valproic acid (3 mM, Acros Organics) were then added to the cells immediately post-transfection to increase recombinant protein production. The cells were harvested 3–5 days after transfection by spinning the cultures at 300g for 5 min. The supernatant was filtered through a 0.45-µm filter and 0.5 mL of 1mM HEPES was added to neutralize the pH. Viral stocks were aliquoted and flash-frozen in liquid nitrogen. They were stored at −80 °C and titrated before further use.

### 2. Supplementary Data

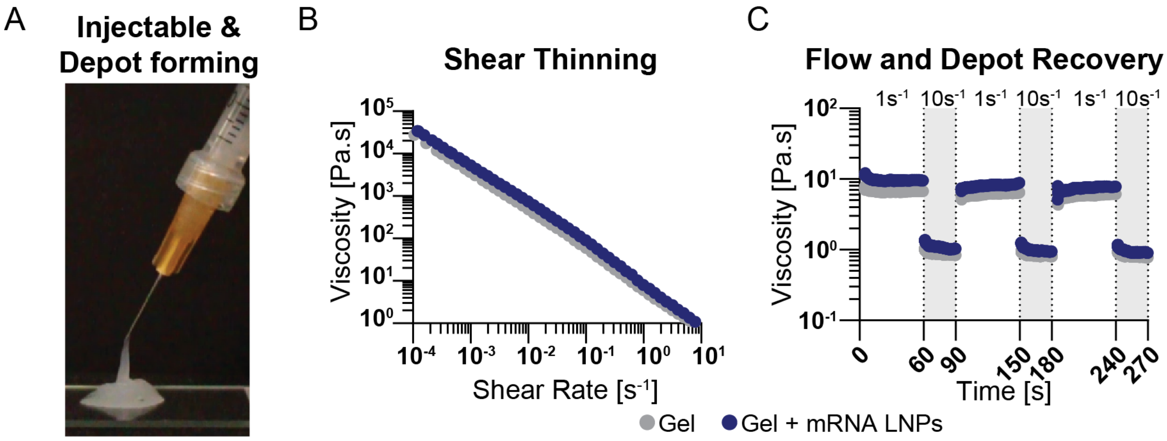


**Fig. S1.**

**PNP hydrogel mechanical characterization.** A) Photo showing PNP-0.5-5 hydrogel is injectable through a 26G needle and forms a solid-like depot following injection. B) High-to-low shear rheology demonstrates shear thinning properties are not impacted by LNP incorporation. C) Step-shear rheology shows PNP hydrogel viscosity drops under high shear (10 $s^{-1}$) and recovers rapidly at low shear (1 $s^{-1}$) over repeated cycles and is not impacted by LNPs.

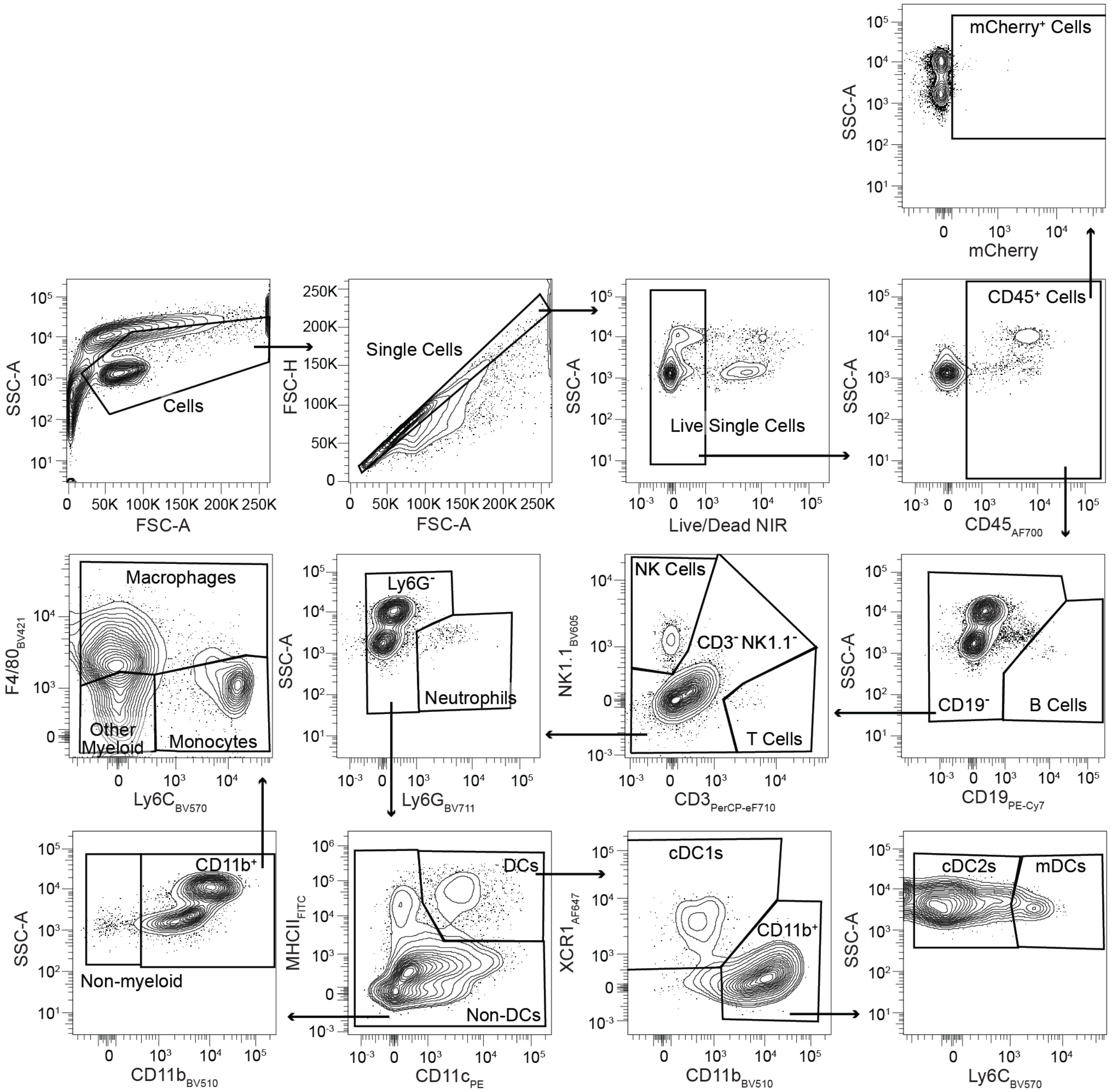


**Fig. S2.**

**PNP hydrogel flow cytometry gating.** Gating strategy for PNP hydrogels shown with representative sample from PNP with 3M-052 on day 3.

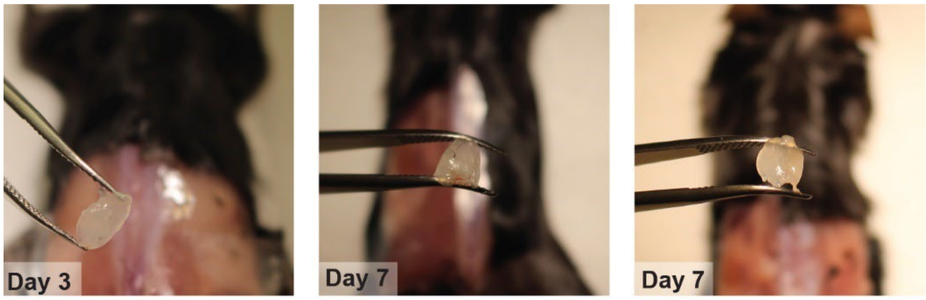


**Fig. S3.**

**PNP hydrogels excised on days 3 and 7.** Photos of PNP-0.5-5 hydrogels on days three and seven show the hydrogel forms a depot without a foreign body fibrotic response and can be easily identified and excised.

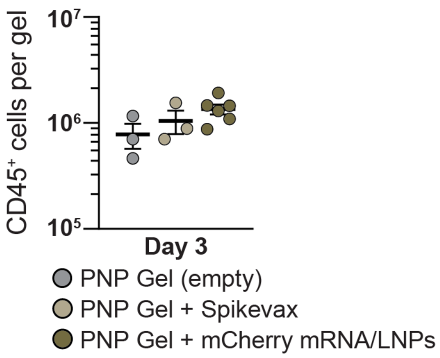


**Fig. S4.**

**PNP hydrogel cell infiltration.** Counts of $CD45^+$ leukocytes on day three in PNP hydrogels with either no cargo (empty), Moderna Spikevax (0.25 µg per dose), or mCherry mRNA/LNPs (1 µg per dose). Data shown as mean ± SEM, n = 3 - 6.

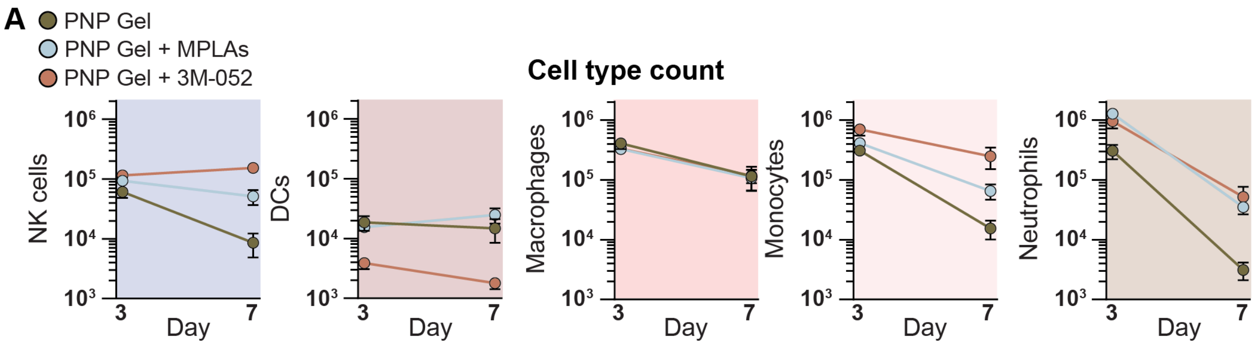


**B** **Cell type % of CD45⁺**

| | Day 3 Gel | Day 3 + MPLAs | Day 3 + 3M-052 | Day 7 Gel | Day 7 + MPLAs | Day 7 + 3M-052 |
|---|---|---|---|---|---|---|
| NK cells | 4.82 | 4.77 | 5.62 | 4.12 | 15.10 | 28.09 |
| T cells | 0.16 | 0.23 | 0.18 | 0.46 | 1.52 | 1.19 |
| B cells | 0.03 | 0.03 | 0.03 | 0.28 | 0.16 | 0.05 |
| DCs | 1.35 | 0.74 | 0.19 | 7.40 | 7.72 | 0.32 |
| Macrophages | 29.77 | 14.82 | 17.06 | 54.58 | 27.25 | 19.33 |
| Monocytes | 23.38 | 18.31 | 32.53 | 9.01 | 22.26 | 36.67 |
| Neutrophils | 24.32 | 58.62 | 41.72 | 2.43 | 13.70 | 7.01 |
| Other myeloid | 14.84 | 1.89 | 1.02 | 18.80 | 9.22 | 3.17 |
| Non-myeloid | 1.32 | 0.58 | 1.66 | 2.90 | 3.06 | 4.17 |

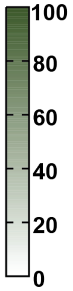


**Fig. S5.**

**PNP hydrogel cell niche.** A) Counts of each cell type per PNP hydrogel depot. Data shown as mean ± SEM, n = 6. B) Percentages of $CD45^+$ gated cells for each cell type. Data shown as mean, n = 6.

A

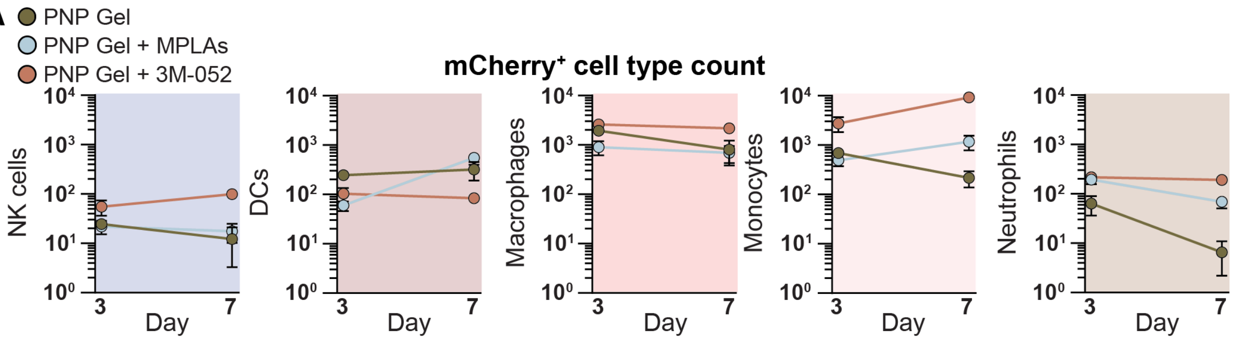


B

**Cell type % of mCherry⁺**

| | Day 3 Gel | Day 3 + MPLAs | Day 3 + 3M-052 | Day 7 Gel | Day 7 + MPLAs | Day 7 + 3M-052 |
|---|---|---|---|---|---|---|
| NK cells | 0.97 | 1.15 | 0.89 | 0.35 | 0.51 | 0.98 |
| T cells | 2.73 | 2.45 | 3.63 | 3.39 | 1.20 | 0.27 |
| B cells | 0.06 | 0.17 | 0.04 | 1.07 | 0.26 | 0.07 |
| DCs | 8.94 | 3.68 | 1.82 | 24.00 | 21.87 | 0.78 |
| Macrophages | 51.31 | 46.38 | 45.23 | 44.36 | 23.07 | 19.26 |
| Monocytes | 18.08 | 29.41 | 42.51 | 19.91 | 48.13 | 75.95 |
| Neutrophils | 3.76 | 11.26 | 3.41 | 0.48 | 3.05 | 1.66 |
| Other myeloid | 13.96 | 5.25 | 2.09 | 5.64 | 1.44 | 0.14 |
| Non-myeloid | 0.20 | 0.25 | 0.38 | 0.81 | 0.47 | 0.89 |

Colour scale: 0, 20, 40, 60, 80, 100

**Fig. S6.**

**mCherry expression in PNP hydrogel cell niche.** A) Counts of mCherry$^{+}$ cells of each cell type per PNP hydrogel depot. Data shown as mean ± SEM, n = 6. B) Percentages of mCherry$^{+}$ CD45$^{+}$ gated cells for each cell type. Data shown as mean, n = 6.

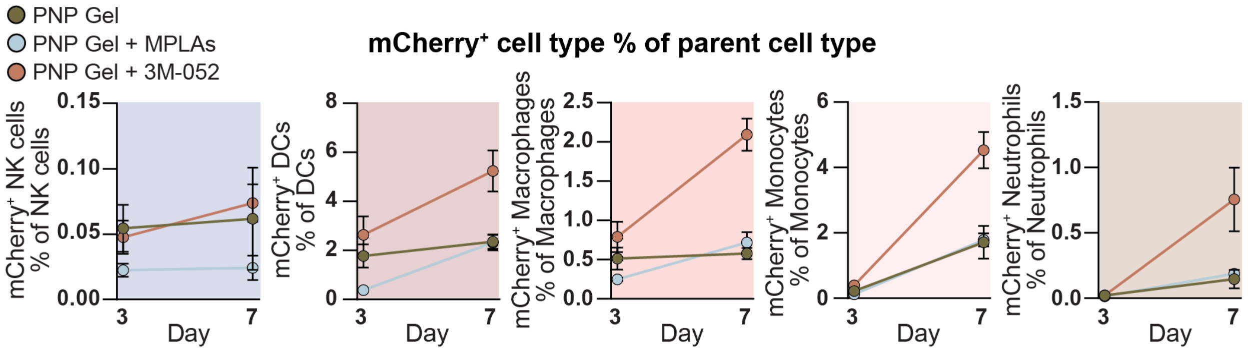


**Fig. S7.**

**mCherry percentage per cell type in PNP hydrogel niche.** Percentage of $mCherry^+$ cells for each cell type. Data shown as mean ± SEM, n = 6.

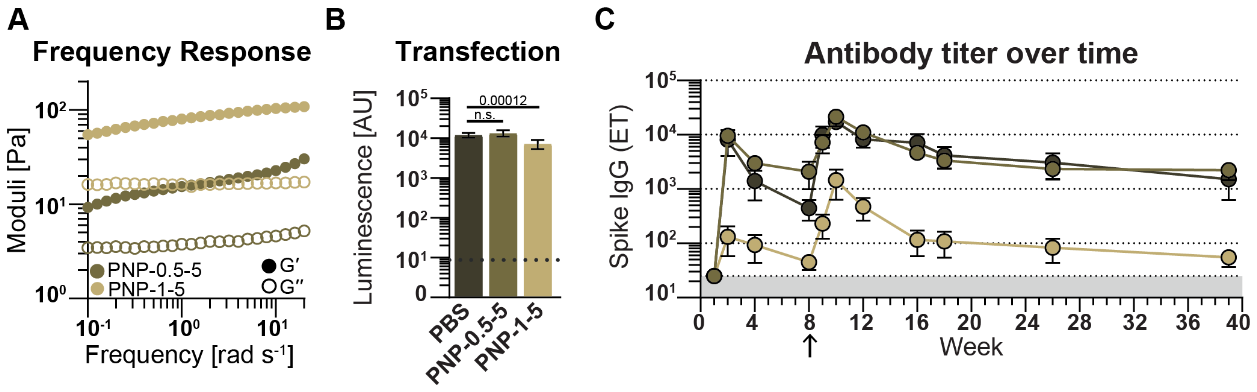


**Fig. S8.**

**Characterization of alternative PNP formulations.** A) Frequency shear rheology shows PNP-1-5 material is stiffer than PNP-0.5-5 and solid-like across time scales. B) Luminescent signal from RAW-Blue macrophages dosed with luciferase mRNA LNPs in either PBS bolus, PNP-0.5-5, or PNP-1-5 hydrogels. Data shown as mean ± SD, n = 8, and statistics are multiple unpaired two-tailed student t-tests run in GraphPad Prism with false discovery rate (FDR) correction using two-stage step-up method of Benjamini, Krieger, and Yekutieli. C) Mice were immunized with 0.25 μg Moderna Spikevax monovalent (WH1) in PBS bolus, PNP-0.5-5, or PNP-1-5 hydrogel at week 0 and boosted with 0.25 μg Moderna Spikevax bivalent in PBS bolus (bolus and PNP-0.5-5 groups) or PNP-1-5 at week eight. Serum was collected and analyzed for anti-spike (WH1) antibodies via ELISA. Data shown as mean ± SEM, n = 5 - 6.

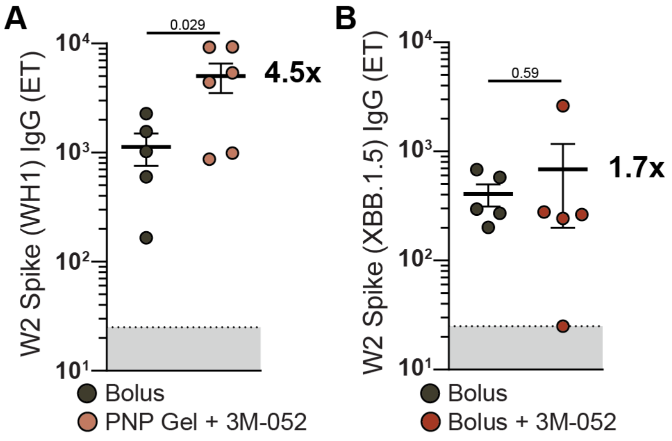


**Fig. S9.**

**Bolus adjuvanting of mRNA vaccine compared with PNP hydrogel and adjuvant.** A) Replicated week two endpoint titer data for bolus control and PNP hydrogel with 3M-052 from figure 4. Statistical values shown are *p* values obtained from GLM fitting and Tukey HSD multiple comparison test in JMP (including groups not plotted here). B) Mice were immunized with 0.25 µg Moderna Spikevax monovalent (XBB.1.5) in PBS bolus with or without 3M-052/Alum. Serum was collected and analyzed for anti-spike (XBB.1.5) antibodies via ELISA. Data shown as mean ± SEM, n = 5, and statistic is an unpaired student t-test run in GraphPad Prism.

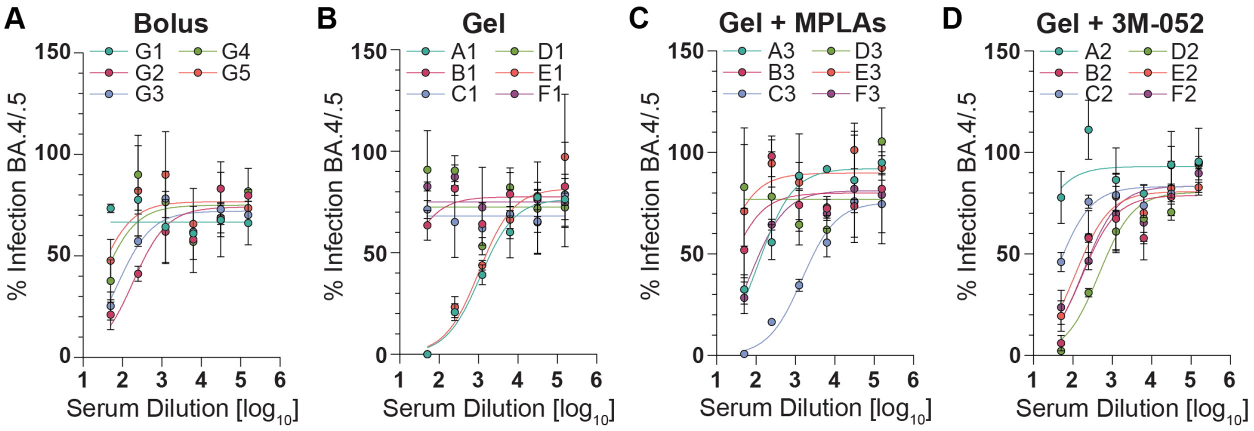


**Fig. S10.**

**Neutralization curves.** Percent infectivity for serial serum dilutions against BA.4/.5 pseudotyped lentivirus for each animal in each treatment group (A-D).

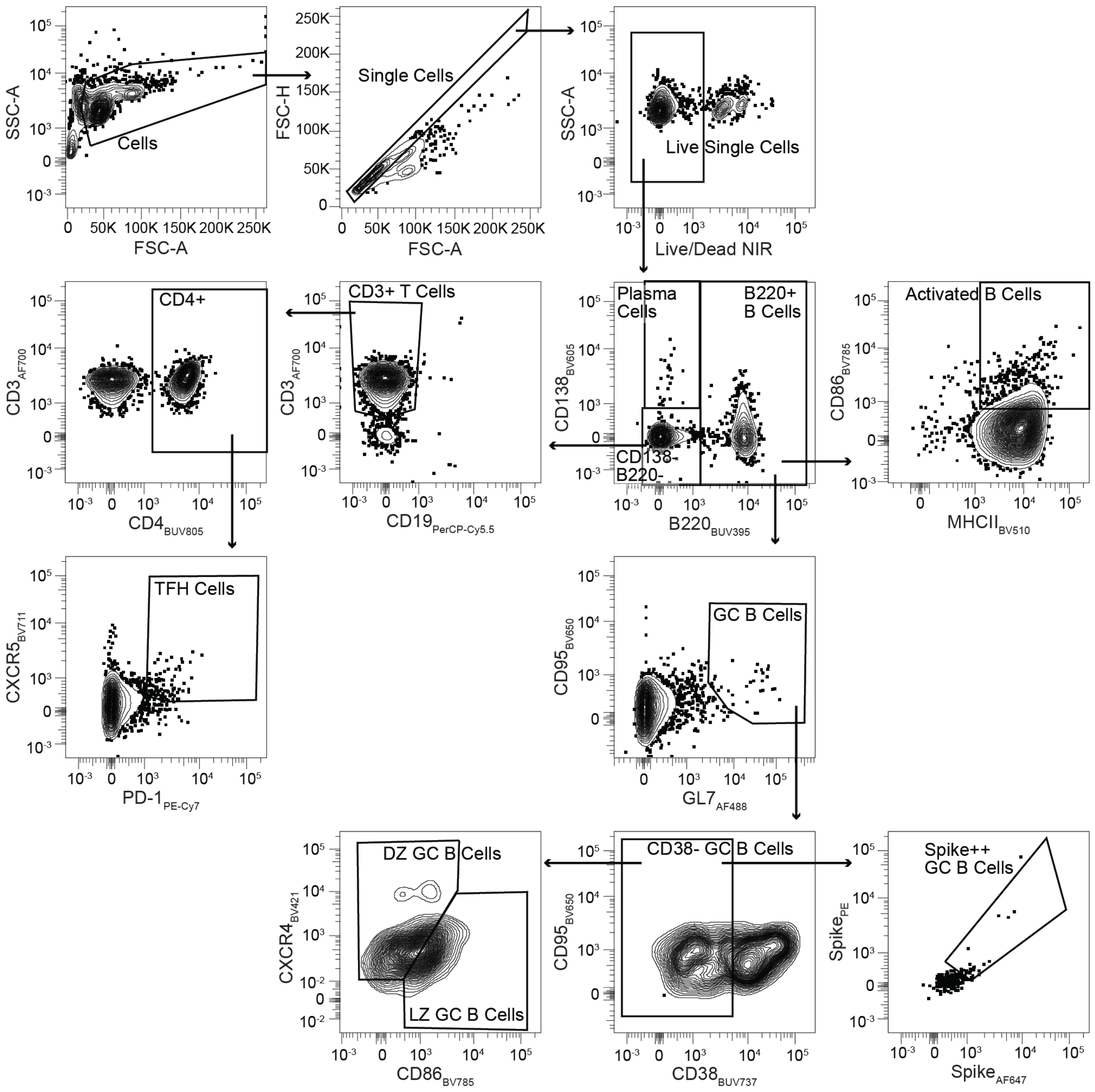


**Fig. S11.**

**Lymph node flow cytometry gating.** Gating strategy for lymph nodes to assess germinal center reactions. Representative sample of PNP hydrogel treatment group on week 1.

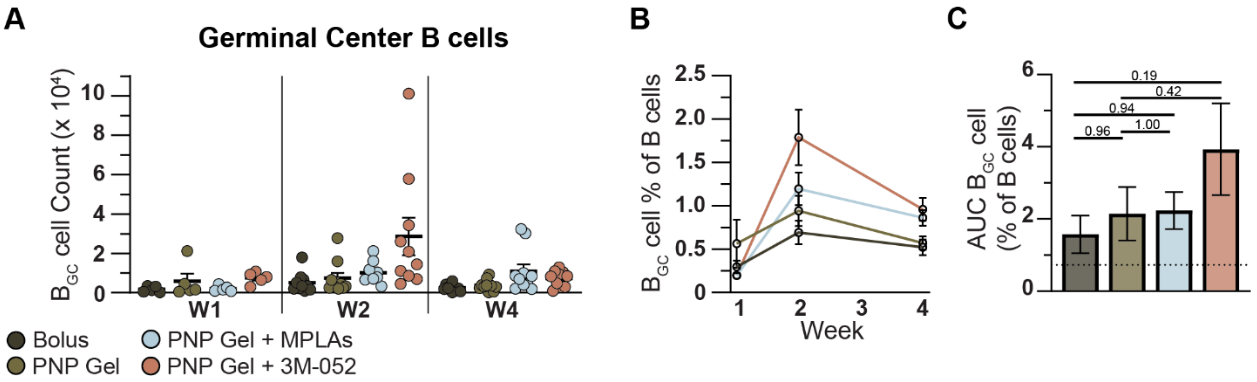


**Fig. S12.**

**Germinal center B cell dynamics** A) Counts of $B_{GC}$ cells. B) $B_{GC}$ cell percentage of B cells over time. Data in A and B shown as mean ± SEM, n = 5 - 10. C) Area under the curve of $B_{GC}$ cell percentage of B cells. Data shown as mean ± SEM. Statistics are one-way ANOVA with Tukey multiple comparisons correction run on data mean, SEM, and N in GraphPad Prism based on recommendations for destructive sampling AUC calculations (https://www.graphpad.com/support/faqid/2031/).

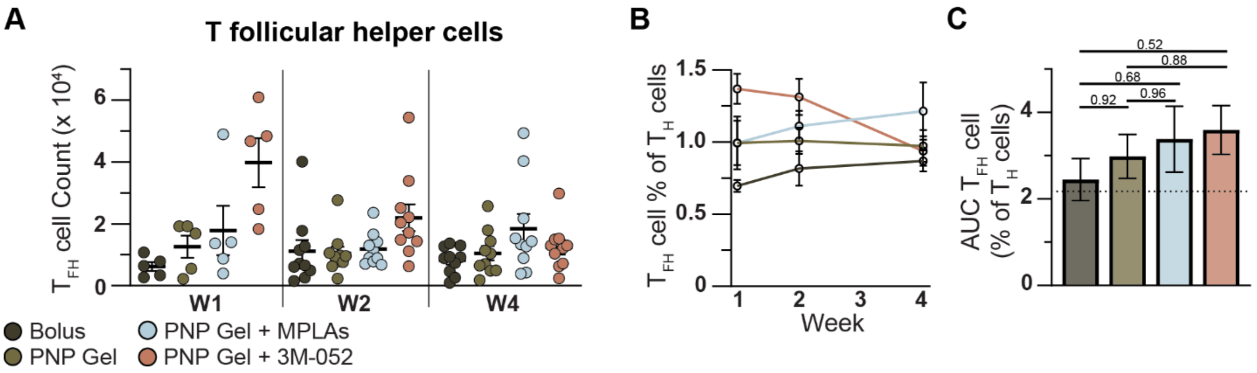


**Fig. S13.**

**Germinal center T follicular helper cell dynamics** Counts of $T_{FH}$ cells. B) $T_{FH}$ cell percentage of $CD4^+$ T cells over time. Data in A and B shown as mean ± SEM, n = 5 - 10. C) Area under the curve of $T_{FH}$ cell percentage of $CD4^+$ T cells. Data shown as mean ± SEM. Statistics are one-way ANOVA with Tukey multiple comparisons correction run on data mean, SEM, and N in GraphPad Prism based on recommendations for destructive sampling AUC calculations.

**Table S1.**

*p* values from general linear model (GLM) followed by Tukey's HSD multiple comparisons procedure comparing among all PNP hydrogel formulations for counts of different infiltrating cell types at days 3 and 7 (referring to Figure S4).

Day 3 cell type counts

| NK cells | | | Adjusted *p* value |
|---|---|---|---|
| PNP gel | vs. | PNP + MPLAs | 0.399 |
| PNP gel | vs. | PNP + 3M-052 | 0.111 |
| PNP + MPLAs | vs. | PNP + 3M-052 | 0.658 |

| Dendritic cells | | | Adjusted *p* value |
|---|---|---|---|
| PNP gel | vs. | PNP + MPLAs | 0.763 |
| PNP gel | vs. | PNP + 3M-052 | **0.021** |
| PNP + MPLAs | vs. | PNP + 3M-052 | 0.066 |

| Macrophages | | | Adjusted *p* value |
|---|---|---|---|
| PNP gel | vs. | PNP + MPLAs | 0.327 |
| PNP gel | vs. | PNP + 3M-052 | 0.366 |
| PNP + MPLAs | vs. | PNP + 3M-052 | 0.996 |

| Monocytes | | | Adjusted *p* value |
|---|---|---|---|
| PNP gel | vs. | PNP + MPLAs | 0.597 |
| PNP gel | vs. | PNP + 3M-052 | **0.010** |
| PNP + MPLAs | vs. | PNP + 3M-052 | **0.049** |

| Neutrophils | | | Adjusted *p* value |
|---|---|---|---|
| PNP gel | vs. | PNP + MPLAs | **0.003** |
| PNP gel | vs. | PNP + 3M-052 | **0.031** |
| PNP + MPLAs | vs. | PNP + 3M-052 | 0.358 |

Day 7 cell type counts

| NK cells | | | Adjusted *p* value |
|---|---|---|---|
| PNP gel | vs. | PNP + MPLAs | 0.122 |
| PNP gel | vs. | PNP + 3M-052 | **< 0.0001** |
| PNP + MPLAs | vs. | PNP + 3M-052 | **0.001** |

| Dendritic cells | | | Adjusted *p* value |
|---|---|---|---|
| PNP gel | vs. | PNP + MPLAs | 0.486 |
| PNP gel | vs. | PNP + 3M-052 | 0.308 |
| PNP + MPLAs | vs. | PNP + 3M-052 | **0.050** |

| Macrophages | | | Adjusted *p* value |
|---|---|---|---|
| PNP gel | vs. | PNP + MPLAs | 0.991 |
| PNP gel | vs. | PNP + 3M-052 | 1.000 |
| PNP + MPLAs | vs. | PNP + 3M-052 | 0.986 |

| Monocytes | | | Adjusted *p* value |
|---|---|---|---|
| PNP gel | vs. | PNP + MPLAs | 0.781 |
| PNP gel | vs. | PNP + 3M-052 | **0.024** |
| PNP + MPLAs | vs. | PNP + 3M-052 | 0.074 |

| Neutrophils | | | Adjusted *p* value |
|---|---|---|---|
| PNP gel | vs. | PNP + MPLAs | 0.295 |
| PNP gel | vs. | PNP + 3M-052 | 0.085 |
| PNP + MPLAs | vs. | PNP + 3M-052 | 0.698 |

**Table S2.**

*p* values from GLM followed by Tukey's HSD multiple comparisons procedure comparing among all PNP hydrogel formulations for counts of mCherry$^+$ cell types on days 3 and 7 (referring to Figure S5).

Day 3 mCherry$^+$ cell type counts

| NK cells | | | Adjusted *p* value |
|---|---|---|---|
| PNP gel | vs. | PNP + MPLAs | 0.985 |
| PNP gel | vs. | PNP + 3M-052 | 0.145 |
| PNP + MPLAs | vs. | PNP + 3M-052 | 0.112 |

| Dendritic cells | | | Adjusted *p* value |
|---|---|---|---|
| PNP gel | vs. | PNP + MPLAs | **0.002** |
| PNP gel | vs. | PNP + 3M-052 | **0.011** |
| PNP + MPLAs | vs. | PNP + 3M-052 | 0.520 |

| Macrophages | | | Adjusted *p* value |
|---|---|---|---|
| PNP gel | vs. | PNP + MPLAs | 0.263 |
| PNP gel | vs. | PNP + 3M-052 | 0.537 |
| PNP + MPLAs | vs. | PNP + 3M-052 | **0.048** |

| Monocytes | | | Adjusted *p* value |
|---|---|---|---|
| PNP gel | vs. | PNP + MPLAs | 0.962 |
| PNP gel | vs. | PNP + 3M-052 | **0.040** |
| PNP + MPLAs | vs. | PNP + 3M-052 | **0.026** |

| Neutrophils | | | Adjusted *p* value |
|---|---|---|---|
| PNP gel | vs. | PNP + MPLAs | **0.043** |
| PNP gel | vs. | PNP + 3M-052 | **0.018** |
| PNP + MPLAs | vs. | PNP + 3M-052 | 0.866 |

Day 7 mCherry$^+$ cell type counts

| NK cells | | | Adjusted *p* value |
|---|---|---|---|
| PNP gel | vs. | PNP + MPLAs | 0.943 |
| PNP gel | vs. | PNP + 3M-052 | **0.001** |
| PNP + MPLAs | vs. | PNP + 3M-052 | **0.002** |

| Dendritic cells | | | Adjusted *p* value |
|---|---|---|---|
| PNP gel | vs. | PNP + MPLAs | 0.390 |
| PNP gel | vs. | PNP + 3M-052 | 0.365 |
| PNP + MPLAs | vs. | PNP + 3M-052 | **0.045** |

| Macrophages | | | Adjusted *p* value |
|---|---|---|---|
| PNP gel | vs. | PNP + MPLAs | 0.967 |
| PNP gel | vs. | PNP + 3M-052 | **0.037** |
| PNP + MPLAs | vs. | PNP + 3M-052 | **0.025** |

| Monocytes | | | Adjusted *p* value |
|---|---|---|---|
| PNP gel | vs. | PNP + MPLAs | 0.804 |
| PNP gel | vs. | PNP + 3M-052 | **0.000** |
| PNP + MPLAs | vs. | PNP + 3M-052 | **0.001** |

| Neutrophils | | | Adjusted *p* value |
|---|---|---|---|
| PNP gel | vs. | PNP + MPLAs | 0.179 |
| PNP gel | vs. | PNP + 3M-052 | **0.001** |
| PNP + MPLAs | vs. | PNP + 3M-052 | **0.009** |

**Table S3.**
*p* values from GLM followed by Tukey's HSD multiple comparisons procedure comparing among all PNP hydrogel formulations for mCherry+ percentage of parent cell type transformed by y=ln(x/(100-x)) to normally distribute data (referring to Figure S6).

Day 3 mCherry$^+$ percent of parent cell type

| NK cells | | | Adjusted *p* value |
|---|---|---|---|
| PNP gel | vs. | PNP + MPLAs | 0.064 |
| PNP gel | vs. | PNP + 3M-052 | 0.936 |
| PNP + MPLAs | vs. | PNP + 3M-052 | 0.112 |

| Dendritic cells | | | Adjusted *p* value |
|---|---|---|---|
| PNP gel | vs. | PNP + MPLAs | **0.003** |
| PNP gel | vs. | PNP + 3M-052 | 0.314 |
| PNP + MPLAs | vs. | PNP + 3M-052 | **0.000** |

| Macrophages | | | Adjusted *p* value |
|---|---|---|---|
| PNP gel | vs. | PNP + MPLAs | 0.174 |
| PNP gel | vs. | PNP + 3M-052 | 0.354 |
| PNP + MPLAs | vs. | PNP + 3M-052 | **0.017** |

| Monocytes | | | Adjusted *p* value |
|---|---|---|---|
| PNP gel | vs. | PNP + MPLAs | 0.204 |
| PNP gel | vs. | PNP + 3M-052 | 0.210 |
| PNP + MPLAs | vs. | PNP + 3M-052 | **0.011** |

| Neutrophils | | | Adjusted *p* value |
|---|---|---|---|
| PNP gel | vs. | PNP + MPLAs | 0.704 |
| PNP gel | vs. | PNP + 3M-052 | 0.765 |
| PNP + MPLAs | vs. | PNP + 3M-052 | 0.323 |

Day 7 mCherry$^+$ percent of parent cell type

| NK cells | | | Adjusted *p* value |
|---|---|---|---|
| PNP gel | vs. | PNP + MPLAs | 0.997 |
| PNP gel | vs. | PNP + 3M-052 | 0.218 |
| PNP + MPLAs | vs. | PNP + 3M-052 | 0.241 |

| Dendritic cells | | | Adjusted *p* value |
|---|---|---|---|
| PNP gel | vs. | PNP + MPLAs | 0.992 |
| PNP gel | vs. | PNP + 3M-052 | **0.014** |
| PNP + MPLAs | vs. | PNP + 3M-052 | **0.011** |

| Macrophages | | | Adjusted *p* value |
|---|---|---|---|
| PNP gel | vs. | PNP + MPLAs | 0.632 |
| PNP gel | vs. | PNP + 3M-052 | **<.0001** |
| PNP + MPLAs | vs. | PNP + 3M-052 | **0.000** |

| Monocytes | | | Adjusted *p* value |
|---|---|---|---|
| PNP gel | vs. | PNP + MPLAs | 0.749 |
| PNP gel | vs. | PNP + 3M-052 | **0.005** |
| PNP + MPLAs | vs. | PNP + 3M-052 | **0.016** |

| Neutrophils | | | Adjusted *p* value |
|---|---|---|---|
| PNP gel | vs. | PNP + MPLAs | 0.221 |
| PNP gel | vs. | PNP + 3M-052 | **0.029** |
| PNP + MPLAs | vs. | PNP + 3M-052 | 0.439 |

**Table S4.**

*p* values from GLM followed by Tukey's HSD multiple comparisons procedure comparing among all vaccine formulations for WH1 spike-specific IgG endpoint titers at several time points (referring to Figure 4B-D).

IgG endpoint titers

| Week 1 | | | Adjusted *p* value |
|---|---|---|---|
| Bolus | vs. | PNP gel | 0.585 |
| Bolus | vs. | PNP + MPLAs | 1.000 |
| Bolus | vs. | PNP + 3M-052 | 0.991 |
| PNP gel | vs. | PNP + MPLAs | 0.548 |
| PNP gel | vs. | PNP + 3M-052 | 0.731 |
| PNP + MPLAs | vs. | PNP + 3M-052 | 0.990 |

| Week 2 | | | Adjusted *p* value |
|---|---|---|---|
| Bolus | vs. | PNP gel | 0.813 |
| Bolus | vs. | PNP + MPLAs | 0.710 |
| Bolus | vs. | PNP + 3M-052 | **0.029** |
| PNP gel | vs. | PNP + MPLAs | 0.997 |
| PNP gel | vs. | PNP + 3M-052 | 0.135 |
| PNP + MPLAs | vs. | PNP + 3M-052 | 0.191 |

| Week 3 | | | Adjusted *p* value |
|---|---|---|---|
| Bolus | vs. | PNP gel | 0.784 |
| Bolus | vs. | PNP + MPLAs | 0.761 |
| Bolus | vs. | PNP + 3M-052 | **0.026** |
| PNP gel | vs. | PNP + MPLAs | 1.000 |
| PNP gel | vs. | PNP + 3M-052 | 0.133 |
| PNP + MPLAs | vs. | PNP + 3M-052 | 0.144 |

| Week 4 | | | Adjusted *p* value |
|---|---|---|---|
| Bolus | vs. | PNP gel | 0.661 |
| Bolus | vs. | PNP + MPLAs | 0.586 |
| Bolus | vs. | PNP + 3M-052 | **0.003** |
| PNP gel | vs. | PNP + MPLAs | 0.999 |
| PNP gel | vs. | PNP + 3M-052 | **0.026** |
| PNP + MPLAs | vs. | PNP + 3M-052 | **0.034** |

| Week 8 | | | Adjusted *p* value |
|---|---|---|---|
| Bolus | vs. | PNP gel | 0.957 |
| Bolus | vs. | PNP + MPLAs | 0.781 |
| Bolus | vs. | PNP + 3M-052 | **0.043** |
| PNP gel | vs. | PNP + MPLAs | 0.966 |
| PNP gel | vs. | PNP + 3M-052 | 0.093 |
| PNP + MPLAs | vs. | PNP + 3M-052 | 0.211 |

| Week 9 | | | Adjusted *p* value |
|---|---|---|---|
| Bolus | vs. | PNP gel | 0.052 |
| Bolus | vs. | PNP + MPLAs | 0.103 |
| Bolus | vs. | PNP + 3M-052 | 0.947 |
| PNP gel | vs. | PNP + MPLAs | 0.982 |
| PNP gel | vs. | PNP + 3M-052 | 0.119 |
| PNP + MPLAs | vs. | PNP + 3M-052 | 0.227 |

| Week 10 | | | Adjusted *p* value |
|---|---|---|---|
| Bolus | vs. | PNP gel | 0.712 |
| Bolus | vs. | PNP + MPLAs | 0.850 |
| Bolus | vs. | PNP + 3M-052 | 0.433 |
| PNP gel | vs. | PNP + MPLAs | 0.993 |
| PNP gel | vs. | PNP + 3M-052 | 0.058 |
| PNP + MPLAs | vs. | PNP + 3M-052 | 0.098 |

| Week 12 | | | Adjusted *p* value |
|---|---|---|---|
| Bolus | vs. | PNP gel | 0.995 |
| Bolus | vs. | PNP + MPLAs | 1.000 |
| Bolus | vs. | PNP + 3M-052 | 0.102 |
| PNP gel | vs. | PNP + MPLAs | 0.994 |
| PNP gel | vs. | PNP + 3M-052 | **0.049** |
| PNP + MPLAs | vs. | PNP + 3M-052 | 0.081 |

| Week 16 | | | Adjusted *p* value |
|---|---|---|---|
| Bolus | vs. | PNP gel | 0.997 |
| Bolus | vs. | PNP + MPLAs | 1.000 |
| Bolus | vs. | PNP + 3M-052 | 0.106 |
| PNP gel | vs. | PNP + MPLAs | 0.990 |
| PNP gel | vs. | PNP + 3M-052 | 0.057 |
| PNP + MPLAs | vs. | PNP + 3M-052 | 0.101 |

| Week 26 | | | Adjusted *p* value |
|---|---|---|---|
| Bolus | vs. | PNP gel | 0.999 |
| Bolus | vs. | PNP + MPLAs | 0.964 |
| Bolus | vs. | PNP + 3M-052 | 0.171 |
| PNP gel | vs. | PNP + MPLAs | 0.907 |
| PNP gel | vs. | PNP + 3M-052 | 0.0996 |
| PNP + MPLAs | vs. | PNP + 3M-052 | 0.266 |

**Table S5.**
*p* values from GLM followed by Tukey's HSD multiple comparisons procedure comparing among all vaccine formulations the area under the curve (AUC) of anti-spike IgG titer from week 1 to 26 (referring to Figure 4E).

IgG endpoint titer AUC week 1-26

| AUC | | | Adjusted *p* value |
|---|---|---|---|
| Bolus | vs. | PNP gel | 0.995 |
| Bolus | vs. | PNP + MPLAs | 0.999 |
| Bolus | vs. | PNP + 3M-052 | 0.071 |
| PNP gel | vs. | PNP + MPLAs | 0.979 |
| PNP gel | vs. | PNP + 3M-052 | **0.033** |
| PNP + MPLAs | vs. | PNP + 3M-052 | 0.072 |

**Table S6.**
*p* values from ordinary one-way ANOVA with Tukey multiple comparisons correction run on data mean, SEM, and N in GraphPad Prism comparing among all vaccine formulations the titer decay half-life from boostrapping procedure (referring to Figure 4F).

| Titer decay half-life | | | |
|---|---|---|---|
| Half-life | | | Adjusted *p* value |
| Bolus | vs. | PNP gel | 0.7658 |
| Bolus | vs. | PNP + MPLAs | **< 0.0001** |
| Bolus | vs. | PNP + 3M-052 | **< 0.0001** |
| PNP gel | vs. | PNP + MPLAs | **< 0.0001** |
| PNP gel | vs. | PNP + 3M-052 | **< 0.0001** |
| PNP + MPLAs | vs. | PNP + 3M-052 | **< 0.0001** |

**Table S7.**

*p* values from GLM followed by Tukey's HSD multiple comparisons procedure comparing among all vaccine formulations for percent infectivity at a 1:50 serum dilution against BA.4/.5 pseudotyped lentivirus neutralization at week 13 (referring to Figure 4G).

Percent infectivity at 1:50 dilution

| Week 13 | | | Adjusted *p* value |
|---|---|---|---|
| Bolus | vs. | PNP gel | 0.9466 |
| Bolus | vs. | PNP + MPLAs | 0.9975 |
| Bolus | vs. | PNP + 3M-052 | 0.924 |
| PNP gel | vs. | PNP + MPLAs | 0.9815 |
| PNP gel | vs. | PNP + 3M-052 | 0.6199 |
| PNP + MPLAs | vs. | PNP + 3M-052 | 0.8303 |

**Table S8.**

*p* values from GLM followed by Tukey's HSD multiple comparisons procedure comparing among all vaccine formulations for WH1 spike-specific IFN-γ producing splenocytes at week 10 (referring to Figure 4H).

Spike-specific IFN-γ splenocytes

| Week 10 | | | Adjusted *p* value |
|---|---|---|---|
| Bolus | vs. | PNP gel | 0.9038 |
| Bolus | vs. | PNP + MPLAs | 0.9967 |
| Bolus | vs. | PNP + 3M-052 | 0.9577 |
| PNP gel | vs. | PNP + MPLAs | 0.2979 |
| PNP gel | vs. | PNP + 3M-052 | 0.6167 |
| PNP + MPLAs | vs. | PNP + 3M-052 | 0.3436 |

**Table S9.**

*p* values from GLM followed by Tukey's HSD multiple comparisons procedure comparing among all vaccine formulations for WH1 spike-specific IgG isotype endpoint titers and ratios at week 16 (referring to Figure 5A, C).

IgG isotype endpoint titer, week 16

| IgG2c | | | Adjusted *p* value |
|---|---|---|---|
| Bolus | vs. | PNP gel | 0.9607 |
| Bolus | vs. | PNP + MPLAs | 0.9807 |
| Bolus | vs. | PNP + 3M-052 | 0.1255 |
| PNP gel | vs. | PNP + MPLAs | 0.9995 |
| PNP gel | vs. | PNP + 3M-052 | **0.037** |
| PNP + MPLAs | vs. | PNP + 3M-052 | **0.0467** |

| IgG1 | | | Adjusted *p* value |
|---|---|---|---|
| Bolus | vs. | PNP gel | 0.6289 |
| Bolus | vs. | PNP + MPLAs | 0.8172 |
| Bolus | vs. | PNP + 3M-052 | 0.9982 |
| PNP gel | vs. | PNP + MPLAs | 0.9849 |
| PNP gel | vs. | PNP + 3M-052 | 0.6999 |
| PNP + MPLAs | vs. | PNP + 3M-052 | 0.8796 |

| IgG2c : IgG1 ratio | | | Adjusted *p* value |
|---|---|---|---|
| Bolus | vs. | PNP gel | 1 |
| Bolus | vs. | PNP + MPLAs | 1 |
| Bolus | vs. | PNP + 3M-052 | 0.5634 |
| PNP gel | vs. | PNP + MPLAs | 1 |
| PNP gel | vs. | PNP + 3M-052 | 0.5248 |
| PNP + MPLAs | vs. | PNP + 3M-052 | 0.5248 |

**Table S10.**

*p* values from GLM followed by Tukey's HSD multiple comparisons procedure comparing among all vaccine formulations for different variant spike-specific IgG endpoint titers at week 16 (referring to Figure 5B).

Variant IgG endpoint titer, week 16

| BA.4/.5 | | | Adjusted *p* value |
|---|---|---|---|
| Bolus | vs. | PNP gel | 1 |
| Bolus | vs. | PNP + MPLAs | 0.8577 |
| Bolus | vs. | PNP + 3M-052 | 0.3098 |
| PNP gel | vs. | PNP + MPLAs | 0.8441 |
| PNP gel | vs. | PNP + 3M-052 | 0.2755 |
| PNP + MPLAs | vs. | PNP + 3M-052 | 0.7259 |

| EG.5.1 | | | Adjusted *p* value |
|---|---|---|---|
| Bolus | vs. | PNP gel | 0.9992 |
| Bolus | vs. | PNP + MPLAs | 0.9934 |
| Bolus | vs. | PNP + 3M-052 | 0.1342 |
| PNP gel | vs. | PNP + MPLAs | 0.9989 |
| PNP gel | vs. | PNP + 3M-052 | 0.1397 |
| PNP + MPLAs | vs. | PNP + 3M-052 | 0.179 |

| B.1.1.529 | | | Adjusted *p* value |
|---|---|---|---|
| Bolus | vs. | PNP gel | 0.9996 |
| Bolus | vs. | PNP + MPLAs | 1 |
| Bolus | vs. | PNP + 3M-052 | 0.3378 |
| PNP gel | vs. | PNP + MPLAs | 0.9999 |
| PNP gel | vs. | PNP + 3M-052 | 0.348 |
| PNP + MPLAs | vs. | PNP + 3M-052 | 0.3146 |

| BA.2.86 | | | Adjusted *p* value |
|---|---|---|---|
| Bolus | vs. | PNP gel | 0.9975 |
| Bolus | vs. | PNP + MPLAs | 0.9999 |
| Bolus | vs. | PNP + 3M-052 | 0.108 |
| PNP gel | vs. | PNP + MPLAs | 0.9993 |
| PNP gel | vs. | PNP + 3M-052 | 0.1255 |
| PNP + MPLAs | vs. | PNP + 3M-052 | 0.0996 |

| XBB.1.5 | | | Adjusted *p* value |
|---|---|---|---|
| Bolus | vs. | PNP gel | 0.9993 |
| Bolus | vs. | PNP + MPLAs | 0.9938 |
| Bolus | vs. | PNP + 3M-052 | 0.1083 |
| PNP gel | vs. | PNP + MPLAs | 0.999 |
| PNP gel | vs. | PNP + 3M-052 | 0.111 |
| PNP + MPLAs | vs. | PNP + 3M-052 | 0.143 |

| SARS-CoV-1 | | | Adjusted *p* value |
|---|---|---|---|
| Bolus | vs. | PNP gel | 0.9999 |
| Bolus | vs. | PNP + MPLAs | 1 |
| Bolus | vs. | PNP + 3M-052 | 0.2107 |
| PNP gel | vs. | PNP + MPLAs | 0.9998 |
| PNP gel | vs. | PNP + 3M-052 | 0.2016 |
| PNP + MPLAs | vs. | PNP + 3M-052 | 0.1777 |

**Table S11.**

*p* values from GLM followed by Tukey's HSD multiple comparisons procedure comparing among all vaccine formulations for variant breadth index at week 16 (referring to Figure 5E).

| Variant breadth index | | | |
|---|---|---|---|
| Week 16 | | | Adjusted *p* value |
| Bolus | vs. | PNP gel | 0.9708 |
| Bolus | vs. | PNP + MPLAs | 0.9638 |
| Bolus | vs. | PNP + 3M-052 | **< 0.0001** |
| PNP gel | vs. | PNP + MPLAs | 1 |
| PNP gel | vs. | PNP + 3M-052 | **< 0.0001** |
| PNP + MPLAs | vs. | PNP + 3M-052 | **< 0.0001** |

**Table S12.**

*p* values from GLM followed by Tukey's HSD multiple comparisons procedure comparing among all vaccine formulations for different variant spike-specific IgG endpoint titer ratios at week 16 (referring to Figure 5F).

Variant IgG endpoint titer ratio to WH1, week 16

| BA.4/.5 | | | Adjusted *p* value |
|---|---|---|---|
| Bolus | vs. | PNP gel | 0.9565 |
| Bolus | vs. | PNP + MPLAs | 0.3897 |
| Bolus | vs. | PNP + 3M-052 | 0.8982 |
| PNP gel | vs. | PNP + MPLAs | 0.6514 |
| PNP gel | vs. | PNP + 3M-052 | 0.9972 |
| PNP + MPLAs | vs. | PNP + 3M-052 | 0.765 |

| EG.5.1 | | | Adjusted *p* value |
|---|---|---|---|
| Bolus | vs. | PNP gel | 0.2097 |
| Bolus | vs. | PNP + MPLAs | 0.4217 |
| Bolus | vs. | PNP + 3M-052 | 0.1551 |
| PNP gel | vs. | PNP + MPLAs | 0.9588 |
| PNP gel | vs. | PNP + 3M-052 | 0.9976 |
| PNP + MPLAs | vs. | PNP + 3M-052 | 0.9024 |

| B.1.1.529 | | | Adjusted *p* value |
|---|---|---|---|
| Bolus | vs. | PNP gel | 0.7242 |
| Bolus | vs. | PNP + MPLAs | 0.9633 |
| Bolus | vs. | PNP + 3M-052 | 0.1517 |
| PNP gel | vs. | PNP + MPLAs | 0.9312 |
| PNP gel | vs. | PNP + 3M-052 | 0.612 |
| PNP + MPLAs | vs. | PNP + 3M-052 | 0.2881 |

| BA.2.86 | | | Adjusted *p* value |
|---|---|---|---|
| Bolus | vs. | PNP gel | 0.729 |
| Bolus | vs. | PNP + MPLAs | 0.9378 |
| Bolus | vs. | PNP + 3M-052 | 0.3691 |
| PNP gel | vs. | PNP + MPLAs | 0.9615 |
| PNP gel | vs. | PNP + 3M-052 | 0.9125 |
| PNP + MPLAs | vs. | PNP + 3M-052 | 0.6703 |

| XBB.1.5 | | | Adjusted *p* value |
|---|---|---|---|
| Bolus | vs. | PNP gel | 0.5254 |
| Bolus | vs. | PNP + MPLAs | 0.7015 |
| Bolus | vs. | PNP + 3M-052 | 0.135 |
| PNP gel | vs. | PNP + MPLAs | 0.9894 |
| PNP gel | vs. | PNP + 3M-052 | 0.7754 |
| PNP + MPLAs | vs. | PNP + 3M-052 | 0.595 |

| SARS-CoV-1 | | | Adjusted *p* value |
|---|---|---|---|
| Bolus | vs. | PNP gel | 0.6826 |
| Bolus | vs. | PNP + MPLAs | 0.9999 |
| Bolus | vs. | PNP + 3M-052 | 0.1846 |
| PNP gel | vs. | PNP + MPLAs | 0.6947 |
| PNP gel | vs. | PNP + 3M-052 | 0.7269 |
| PNP + MPLAs | vs. | PNP + 3M-052 | 0.176 |

**Table S13.**

*p* values from GLM followed by Tukey's HSD multiple comparisons procedure comparing among all vaccine formulations for activated B cell percentage of all B cells at week one, transformed by y=ln(x/(100-x)) to normally distribute data (referring to Figure 6B).

Activated B cell percentage of B cells

| Week 1 | | | Adjusted *p* value |
|---|---|---|---|
| Bolus | vs. | PNP gel | 0.4118 |
| Bolus | vs. | PNP + MPLAs | 0.1726 |
| Bolus | vs. | PNP + 3M-052 | **0.0003** |
| PNP gel | vs. | PNP + MPLAs | 0.925 |
| PNP gel | vs. | PNP + 3M-052 | **0.0037** |
| PNP + MPLAs | vs. | PNP + 3M-052 | **0.0108** |

**Table S14.**

*p* values from GLM followed by Tukey's HSD multiple comparisons procedure comparing among all vaccine formulations for germinal center B ($B_{GC}$) cell light zone : dark zone ratio (LZ:DZ) at week one (referring to Figure 6C).

$B_{GC}$ cell LZ:DZ ratio

| Week 1 | | | Adjusted *p* value |
|---|---|---|---|
| Bolus | vs. | PNP gel | 0.0899 |
| Bolus | vs. | PNP + MPLAs | 0.7119 |
| Bolus | vs. | PNP + 3M-052 | **0.0003** |
| PNP gel | vs. | PNP + MPLAs | 0.4397 |
| PNP gel | vs. | PNP + 3M-052 | **0.0246** |
| PNP + MPLAs | vs. | PNP + 3M-052 | **0.0017** |

**Table S15.**

*p* values from GLM followed by Tukey's HSD multiple comparisons procedure comparing among all vaccine formulations for $B_{GC}$ cells as a percentage of all B cells across different weeks, transformed by y=ln(x/(100-x)) to normally distribute data (referring to Figure 6G).

$B_{GC}$ cells percent of all B cells

| Week 1 | | | Adjusted *p* value |
|---|---|---|---|
| Bolus | vs. | PNP gel | 0.7861 |
| Bolus | vs. | PNP + MPLAs | 0.8759 |
| Bolus | vs. | PNP + 3M-052 | 0.8643 |
| PNP gel | vs. | PNP + MPLAs | 0.3725 |
| PNP gel | vs. | PNP + 3M-052 | 0.3597 |
| PNP + MPLAs | vs. | PNP + 3M-052 | 1 |
| **Week 2** | | | **Adjusted *p* value** |
| Bolus | vs. | PNP gel | 0.6925 |
| Bolus | vs. | PNP + MPLAs | 0.1415 |
| Bolus | vs. | PNP + 3M-052 | **0.0066** |
| PNP gel | vs. | PNP + MPLAs | 0.6765 |
| PNP gel | vs. | PNP + 3M-052 | 0.0835 |
| PNP + MPLAs | vs. | PNP + 3M-052 | 0.5295 |
| **Week 4** | | | **Adjusted *p* value** |
| Bolus | vs. | PNP gel | 0.9479 |
| Bolus | vs. | PNP + MPLAs | **0.0498** |
| Bolus | vs. | PNP + 3M-052 | **0.0267** |
| PNP gel | vs. | PNP + MPLAs | 0.1499 |
| PNP gel | vs. | PNP + 3M-052 | 0.0871 |
| PNP + MPLAs | vs. | PNP + 3M-052 | 0.9924 |

**Table S16.**

*p* values from GLM followed by Tukey's HSD multiple comparisons procedure comparing among all vaccine formulations for counts of spike-specific (spike$^{++}$) $B_{GC}$ cells across different weeks (referring to Figure 6H).

Spike specific $B_{GC}$ cell counts

| Week 1 | | | Adjusted *p* value |
|---|---|---|---|
| Bolus | vs. | PNP gel | 0.839 |
| Bolus | vs. | PNP + MPLAs | 0.8743 |
| Bolus | vs. | PNP + 3M-052 | **0.0335** |
| PNP gel | vs. | PNP + MPLAs | 0.9998 |
| PNP gel | vs. | PNP + 3M-052 | 0.1367 |
| PNP + MPLAs | vs. | PNP + 3M-052 | 0.1203 |

| Week 2 | | | Adjusted *p* value |
|---|---|---|---|
| Bolus | vs. | PNP gel | 0.9435 |
| Bolus | vs. | PNP + MPLAs | 0.9978 |
| Bolus | vs. | PNP + 3M-052 | 0.2945 |
| PNP gel | vs. | PNP + MPLAs | 0.8804 |
| PNP gel | vs. | PNP + 3M-052 | 0.6054 |
| PNP + MPLAs | vs. | PNP + 3M-052 | 0.2193 |

| Week 4 | | | Adjusted *p* value |
|---|---|---|---|
| Bolus | vs. | PNP gel | 0.8253 |
| Bolus | vs. | PNP + MPLAs | **0.0285** |
| Bolus | vs. | PNP + 3M-052 | 0.8163 |
| PNP gel | vs. | PNP + MPLAs | 0.17 |
| PNP gel | vs. | PNP + 3M-052 | 1 |
| PNP + MPLAs | vs. | PNP + 3M-052 | 0.1756 |

**Table S17.**

*p* values from GLM followed by Tukey's HSD multiple comparisons procedure comparing among all vaccine formulations for T follicular helper ($T_{FH}$) cells as a percentage of all $CD4^+$ T cells across different weeks, transformed by y=ln(x/(100-x)) to normally distribute data (referring to Figure 6I).

$T_{FH}$ percent of $CD4^+$ T cells

| Week 1 | | | Adjusted *p* value |
|---|---|---|---|
| Bolus | vs. | PNP gel | 0.2741 |
| Bolus | vs. | PNP + MPLAs | 0.3037 |
| Bolus | vs. | PNP + 3M-052 | **0.0073** |
| PNP gel | vs. | PNP + MPLAs | 0.9998 |
| PNP gel | vs. | PNP + 3M-052 | 0.19 |
| PNP + MPLAs | vs. | PNP + 3M-052 | 0.1696 |

| Week 2 | | | Adjusted *p* value |
|---|---|---|---|
| Bolus | vs. | PNP gel | 0.2584 |
| Bolus | vs. | PNP + MPLAs | 0.0628 |
| Bolus | vs. | PNP + 3M-052 | **0.0032** |
| PNP gel | vs. | PNP + MPLAs | 0.8771 |
| PNP gel | vs. | PNP + 3M-052 | 0.2159 |
| PNP + MPLAs | vs. | PNP + 3M-052 | 0.6047 |

| Week 4 | | | Adjusted *p* value |
|---|---|---|---|
| Bolus | vs. | PNP gel | 0.9606 |
| Bolus | vs. | PNP + MPLAs | 0.3468 |
| Bolus | vs. | PNP + 3M-052 | 0.9851 |
| PNP gel | vs. | PNP + MPLAs | 0.6306 |
| PNP gel | vs. | PNP + 3M-052 | 0.999 |
| PNP + MPLAs | vs. | PNP + 3M-052 | 0.5442 |

**Table S18.**

*p* values from GLM followed by Tukey's HSD multiple comparisons procedure comparing among all vaccine formulations for $B_{GC}$ cell ratio to $T_{FH}$ cells at weeks two and four (referring to Figure 6J).

$B_{GC}$ : $T_{FH}$ cell ratio

| Week 2 | | | Adjusted *p* value |
|---|---|---|---|
| Bolus | vs. | PNP gel | 0.9979 |
| Bolus | vs. | PNP + MPLAs | 0.406 |
| Bolus | vs. | PNP + 3M-052 | 0.1452 |
| PNP gel | vs. | PNP + MPLAs | 0.5097 |
| PNP gel | vs. | PNP + 3M-052 | 0.201 |
| PNP + MPLAs | vs. | PNP + 3M-052 | 0.9206 |

| Week 4 | | | Adjusted *p* value |
|---|---|---|---|
| Bolus | vs. | PNP gel | 0.98 |
| Bolus | vs. | PNP + MPLAs | 0.1009 |
| Bolus | vs. | PNP + 3M-052 | **0.019** |
| PNP gel | vs. | PNP + MPLAs | 0.2074 |
| PNP gel | vs. | PNP + 3M-052 | **0.0459** |
| PNP + MPLAs | vs. | PNP + 3M-052 | 0.8715 |

**Table S19.**

*p* values from GLM followed by Tukey's HSD multiple comparisons procedure comparing among all vaccine formulations for counts of $B_{GC}$ cells across different weeks (referring to Figure S11A).

$B_{GC}$ cell counts

| Week 1 | | | Adjusted *p* value |
|---|---|---|---|
| Bolus | vs. | PNP gel | 0.5452 |
| Bolus | vs. | PNP + MPLAs | 0.9992 |
| Bolus | vs. | PNP + 3M-052 | 0.2774 |
| PNP gel | vs. | PNP + MPLAs | 0.6195 |
| PNP gel | vs. | PNP + 3M-052 | 0.9469 |
| PNP + MPLAs | vs. | PNP + 3M-052 | 0.331 |

| Week 2 | | | Adjusted *p* value |
|---|---|---|---|
| Bolus | vs. | PNP gel | 0.9842 |
| Bolus | vs. | PNP + MPLAs | 0.8561 |
| Bolus | vs. | PNP + 3M-052 | **0.005** |
| PNP gel | vs. | PNP + MPLAs | 0.9712 |
| PNP gel | vs. | PNP + 3M-052 | **0.0121** |
| PNP + MPLAs | vs. | PNP + 3M-052 | **0.0341** |

| Week 4 | | | Adjusted *p* value |
|---|---|---|---|
| Bolus | vs. | PNP gel | 0.9634 |
| Bolus | vs. | PNP + MPLAs | **0.019** |
| Bolus | vs. | PNP + 3M-052 | 0.3013 |
| PNP gel | vs. | PNP + MPLAs | 0.056 |
| PNP gel | vs. | PNP + 3M-052 | 0.5651 |
| PNP + MPLAs | vs. | PNP + 3M-052 | 0.5253 |

**Table S20.**

*p* values from one-way ANOVA with Tukey multiple comparisons correction run on data mean, SEM, and N in GraphPad Prism (based on recommendations for destructive sampling AUC calculations, https://www.graphpad.com/support/faqid/2031/) comparing among all vaccine formulations for AUC of $B_{GC}$ cell percentage of all B cells from week one to four (referring to Figure S11C).

$B_{GC}$ cell percentage of all B cells AUC

| AUC | | | Adjusted *p* value |
|---|---|---|---|
| Bolus | vs. | PNP gel | 0.9615 |
| Bolus | vs. | PNP + MPLAs | 0.9416 |
| Bolus | vs. | PNP + 3M-052 | 0.1865 |
| PNP gel | vs. | PNP + MPLAs | 0.9998 |
| PNP gel | vs. | PNP + 3M-052 | 0.42 |
| PNP + MPLAs | vs. | PNP + 3M-052 | 0.4666 |

**Table S21.**
*p* values from GLM followed by Tukey's HSD multiple comparisons procedure comparing among all vaccine formulations for counts of $T_{FH}$ cells across different weeks (referring to Figure S12A).

$T_{FH}$ cell counts

| Week 1 | | | Adjusted *p* value |
|---|---|---|---|
| Bolus | vs. | PNP gel | 0.8211 |
| Bolus | vs. | PNP + MPLAs | 0.4246 |
| Bolus | vs. | PNP + 3M-052 | **0.0032** |
| PNP gel | vs. | PNP + MPLAs | 0.8902 |
| PNP gel | vs. | PNP + 3M-052 | **0.0143** |
| PNP + MPLAs | vs. | PNP + 3M-052 | **0.0497** |

| Week 2 | | | Adjusted *p* value |
|---|---|---|---|
| Bolus | vs. | PNP gel | 0.9973 |
| Bolus | vs. | PNP + MPLAs | 0.9964 |
| Bolus | vs. | PNP + 3M-052 | **0.019** |
| PNP gel | vs. | PNP + MPLAs | 0.976 |
| PNP gel | vs. | PNP + 3M-052 | **0.0119** |
| PNP + MPLAs | vs. | PNP + 3M-052 | **0.0314** |

| Week 4 | | | Adjusted *p* value |
|---|---|---|---|
| Bolus | vs. | PNP gel | 0.9176 |
| Bolus | vs. | PNP + MPLAs | **0.0348** |
| Bolus | vs. | PNP + 3M-052 | 0.6346 |
| PNP gel | vs. | PNP + MPLAs | 0.1334 |
| PNP gel | vs. | PNP + 3M-052 | 0.9445 |
| PNP + MPLAs | vs. | PNP + 3M-052 | 0.3415 |

**Table S22.**

*p* values from one-way ANOVA with Tukey multiple comparisons correction run on data mean, SEM, and N in GraphPad Prism comparing among all vaccine formulations for area under the curve of $T_{FH}$ cell percentage of all CD4+ T cells from week one to four (referring to Figure S12C).

$T_{FH}$ percentage of $CD4^+$ T cell AUC

| AUC | | | Adjusted *p* value |
|---|---|---|---|
| Bolus | vs. | PNP gel | 0.9171 |
| Bolus | vs. | PNP + MPLAs | 0.6766 |
| Bolus | vs. | PNP + 3M-052 | 0.5184 |
| PNP gel | vs. | PNP + MPLAs | 0.9638 |
| PNP gel | vs. | PNP + 3M-052 | 0.8848 |
| PNP + MPLAs | vs. | PNP + 3M-052 | 0.9943 |
